# Spatially ordered zygotic genome activation fulfills embryo quality control

**DOI:** 10.1101/2024.12.22.629969

**Authors:** Wenchao Qian, Hui Chen, Hongju Lee, Matthew C. Good

**Affiliations:** Department of Cell and Developmental Biology, Perelman School of Medicine, University of Pennsylvania, Philadelphia, PA 19104, USA; Cell and Molecular Biology Graduate Group, Perelman School of Medicine, University of Pennsylvania, Philadelphia, PA 19104, USA; Department of Bioengineering, School of Engineering and Applied Science, University of Pennsylvania, Philadelphia, PA 19104, USA

**Keywords:** zygotic genome activation, spatiotemporal ZGA, cell size, blastula patterning, quality control, maternal-to-zygotic transition, embryo temperature controller, cell division

## Abstract

Early embryo development features autonomous, maternally-driven cell divisions that self-organize the multicellular blastula or blastocyst tissue. Maternal control cedes to the zygote starting with the onset of widespread zygotic genome activation (ZGA), which is essential for subsequent cell fate determination and morphogenesis. Intriguingly, although the onset of ZGA is highly regulated at the level of an embryo, it can be non-homogenous and precisely patterned at the single-cell level. We previously demonstrated a stereotyped spatial and temporal ordering of ZGA in a model vertebrate embryo. Unknown, however, was whether this precise ZGA patterning was required for development. To address this fundamental question, we devised a strategy to spatially control cell divisions in the embryo that perturb blastula embryo organization. We demonstrate the feasibility of spatially inverting the cell size pattern of embryos and find that these inverted embryos undergo a flipped pattern of ZGA. Mispatterned ZGA along the animal-vegetal axis causes embryo apoptosis, revealing that gastrula embryos have a built-in quality control system to sense inappropriate ZGA patterning, including regional defects in transcriptional onset. The quality control response is non-autonomous which may depend on anti-apoptotic signals that repress cell death outside of the animal hemisphere. These results reveal the requirement of properly patterned ZGA for normal development and the existence of an embryo quality control response exquisitely tuned to the spatial and temporal ordering of genome activation and zygotic gene expression.

## Introduction

The early cleavage stages of embryo development are governed by maternal RNAs deposited in the egg and by translated proteins following fertilization. At a defined interval, the embryo begins the maternal-to-zygotic transition and hands off developmental control from maternal factors to those expressed by cells of the zygote [1–4]. This transition requires the onset of embryonic transcription within nuclei in the process of zygotic genome activation [5–9]. This is an essential developmental event, without which embryos cannot initiate cell fate determination or morphogenesis [3, 10–12]. The timing of ZGA onset differs between species [6–8, 13], but within an organism, it is highly stereotyped and reproducible. Numerous control systems have been described, and chromatin opening and transcription factors binding are required for transcription onset in *Drosophila*, zebrafish, *Xenopus*, and mammalian embryos [14–21]. In non-placental embryo models, the onset of ZGA is not dictated by elapsed time post-fertilization but is instead coupled to reaching a critical cell size (cell diameter) or DNA-to-cytoplasm ratio (DC ratio) [2, 3, 22]. Alterations of cell size or DC ratio, also classically termed nucleocytoplasmic ratio (NC ratio), are sufficient to alter the timing of large-scale ZGA [3, 23–26].

Intriguingly, the onset of ZGA is not spatially uniform in a variety of model embryonic systems [23, 27]. For example, the *Xenopus* blastula displays a highly stereotyped spatiotemporal pattern of ZGA onset, that is coupled to an animal-vegetal (A-V) cell size gradient [23]. At the 8- cell stage, the embryo contains animal and vegetal blastomeres. Although all cells generally divide quite quickly, the blastomeres in the AP divide more rapidly [28] and therefore are smaller in size than those of the VP at the onset of the major wave of ZGA [23]. We have theorized that spatial and temporal ordering of ZGA onset may be linked to the temporal ordering of cell fate determination [29]. However, to date, the function of spatially graded ZGA has remained a mystery.

Revealing an essential development role for patterned ZGA onset requires perturbation of the endogenous blastula tissue organization to alter the spatial order or transcriptional onset. Precise spatial manipulation of the cell division period in the zygote can be achieved using embryo temperature controllers. Previous efforts to manipulate the cellular organization of non-placental embryos have leveraged localized thermal control of cell division periods, including via focused heating, or by housing embryos in microfluidic devices or within microchambers [28, 30–33]. Such a device was developed for millimeter-sized *Xenopus* embryos and has been shown capable of spatially controlling blastomere cell division rates [28].

In this study, we sought to further develop this embryo temperature controller to investigate how blastula tissue organization controls ZGA patterning and subsequent zygote development. We demonstrate successful spatial control of the cell division period to reconfigure cell size gradients in the blastula, including inversion of the endogenous A-V cell size gradient. This inverted blastula shows a flipped pattern of ZGA and inverted spatiotemporal onset of zygotic transcription: vegetal-first rather than animal-first. We determine that proper spatial patterning of ZGA onset is essential for normal development. Mispatterned ZGA leads to activation of embryo apoptosis, dose-dependent based on the extent of ZGA delay in the animal pole. Additionally, we determine that gastrula embryos have a built-in quality control system that senses inappropriate ZGA onset, including delay of ZGA in a subregion of embryos. The quality control response is non-autonomous, as cells that die first are not necessarily those delayed in ZGA onset. These results provide direct evidence that spatial and temporal ordering of ZGA is essential for early development and reveal a previously uncharacterized quality control response within embryos capable of sensing mispatterned ZGA.

## Results

### Controlling the spatial pattern of cell divisions in the blastula

To determine the biological role of spatiotemporally ordered ZGA onset, we sought to alter blastula tissue organization. The speed of cell division of non-placental embryos is dependent on temperature [34] (**Fig. S1A**). *Xenopus* embryos tolerate a range of temperatures, and their cell division rate speeds up when above room temperature and slows down when below it. Various groups have controlled cell division speed in spatial subregions of the embryo by application of temperature gradients [28, 31–33]. A device was specifically devised for millimeter size *Xenopus* embryos, which are placed between two thin aluminum plates (**Fig. S1B-D**). The temperature of each plate is controlled by peltiers, that enable the creation of a temperature gradient to spatially regulate the internal cell division period in the cleavage stage blastula embryo [28]. We sought to further develop and expand this embryo temperature controller to investigate how blastula tissue organization controls ZGA patterning and zygote development.

To invert the pattern of the cell division period along the A-V axis, we cooled the plate proximal to the normally fast-dividing animal pole cells and warmed the plate next to the normally slower-dividing vegetal pole cells. Based on a similar device, we expected this would slow down the cell cycle period in animal cells and speed up the divisions in vegetal cells [28]. Importantly, embryos are only inside the chamber during cleavage stage divisions of blastula development (2.5 – 9 hpf, **Fig. 1A, S1B-D**) prior to any morphogenesis. Thus, only cell division speeds and cell size gradient are altered within the blastula, and embryos are then removed from the device and transferred back to dishes containing media at room temperature for subsequent gastrulation and later development.

**Figure 1.**
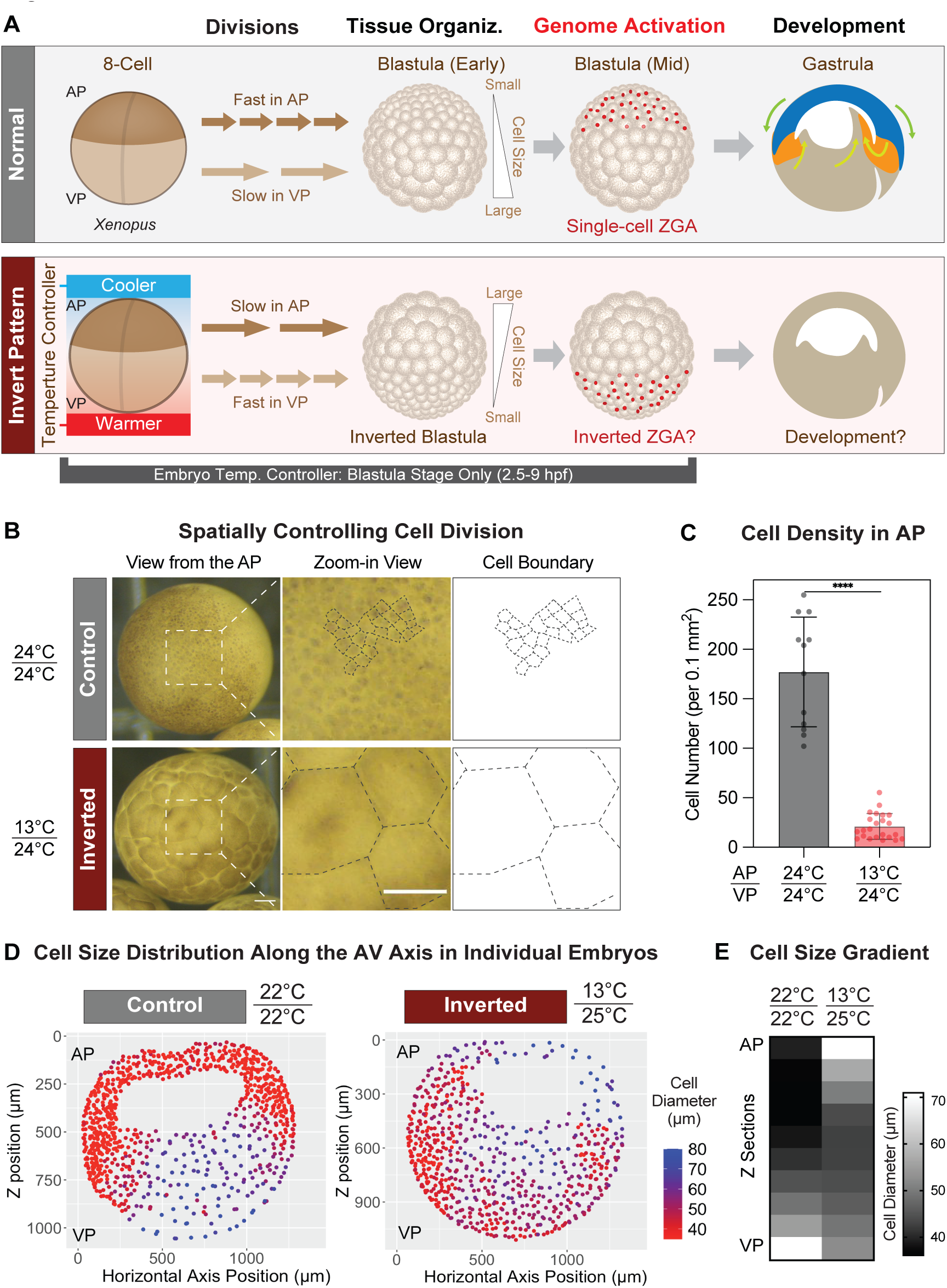
Spatial control of cell division speed within the embryo enables inversion of blastula cell size patterning. **(A)** Schematic showing regionalized division speed, embryo blastula cell size, and subsequent development in a *Xenopus* embryo (top row). Incubation of a zygote within embryo temperature controller (bottom row) (blastula stage: 2.5-9 hpf) enables spatial control of cell division period. Cooling cells within the AP and warming cells in the VP is predicted to invert blastula cell size patterning. Unknown is whether this will invert ZGA patterning and impact embryo development. AP, animal pole. VP, vegetal pole. **(B)** Representative stereomicroscope images showing cells of the animal pole for late-blastula embryos (9.2 hpf) that were incubated in chambers: 24°C/24°C control and 13°C/24°C (AP temp/ VP temp). Zoom-in with cell boundaries. Scale bar, 0.1 mm. **(C)** Average cell number per area (0.1 mm^2^) on the animal pole surface for control 24/24°C and temperature gradient embryos at 9.2 hpf. Mean +- SD. N=12 embryos for 24/24°C control, N=23 embryos for 13°C/24°C. Mann-Whitney test, P<0.0001. **(D)** Sagittal slice of 3D reconstructed embryo fixed at 9.5 hpf, showing cell diameter (nucleus-to- nucleus distance) as a heatmap. Individual nuclei color-coded based on cell diameter. Red: smallest; blue largest. Cell size gradient is inverted within 13°C/25°C Shows nuclei from center section (1/10^th^ slice). **(E)** Average cell diameter as a function of position along animal-vegetal (AV) axis at 9.5 hpf. Mean diameter for cells within each of 10 sections along AV axis. N=2 embryos for 22°C/22°C, N=6 embryos for 13°C/24°C. Shows inversion of cell size gradient.

We hypothesized that the spatial inversion of the cell division period would result in an inverted pattern of cell sizes, such that in the late blastula, animal cells would be larger than the vegetal cells (**Fig. 1A**). After optimization, we settled on a temperature gradient of 13°C toward AP cells and 24°C toward VP cells. In this configuration, we found that late blastula embryos had extremely large cell sizes on the surface of the animal pole compared to control embryos in chambers set to 24°C on both plates (**Fig. 1B**). The cell number in the entire AP was greatly reduced by more than 11-fold (**Fig. 1C**). In contrast, the VP cells were largely the same size (**Fig S1H**), as expected. These data indicate a successful slowing of AP cell division speed.

To determine the overall cellular organization of the blastula, we performed whole-mount DNA staining of embryos in temperature gradient versus control chambers. To quantify cell diameter for these roughly spheroid blastomeres we quantified average inter-nuclei distance as a proxy, as has been done previously [23]. A sagittal section view of the embryo showing the A-V axis reveals the extent of alteration to late blastula organization in embryos developed within a temperature gradient versus a uniform temperature. The blastula develops normally and creates a fluid-filled blastocoel cavity. However, the cell size gradient along the A-V axis is inverted (**Fig. 1D, E**). In control embryos (developed in a uniform 24°C chamber), the smallest cells are primarily located in the animal cap, and the largest cells are found in the vegetal mass. In contrast, in embryos incubated in a temperature gradient, the largest cells were now present in the animal cap, and the smallest cells were found in deep vegetal regions (**Fig. 1D**). Averaging cell sizes in bins along the A-V axis further reveals an inversion of the cell size gradient (**Fig 1E**). Together, these results demonstrate that our temperature gradient was largely effective at inverting blastula cell size organization. We refer to these temperature gradient embryos ‘inverted embryos’ moving forward in the study.

We next asked whether the temperature gradient within embryos was close to linear and whether embryos retain a “memory” of the device once removed from it. It is not technically feasible to insert a temperature probe or thermistor within the embryo. Instead, we converted the cell density of each vertical section to extrapolate the number of cell divisions that had occurred and the effective temperature of each section using data from embryos that grew in uniform temperatures (either 13°C or 24°C). We found that the temperature gradient along the AV is close to linear (**Fig. S1E**), confirming how blastula inversion was achieved. Additionally, we asked whether the inverted embryos resume normal divisions after being removed from the chambers at 9 hpf. Cleavage stage divisions at room temperature are rapid in the AP with an average period of 27 minutes at 23°C (**Fig. S1F**). The cell cycle matures and extends at the mid-blastula transition when cells reach a threshold cell size [1, 35]. If blastomeres had a memory of the device, they would continue to divide at a temperature-controlled rate within the device. Alternatively, if development proceeded normally (and without memory), we expected AP cells would resume very rapid divisions at 23°C. We found that AP cells of the inverted embryos quickly resumed fast division rates (∼ 2 divisions/hour), and did not continue the slow divisions that occurred under the temperature gradients (**Fig. S1G** and **Video 1**). This is likely due to their large sizes and not having reached the cell size threshold for cell cycle elongation. In contrast, AP cells in control embryos were already very small and had elongated the cell cycle period. Thus, cells the inverted embryos do not have a memory of the device. Instead, they continue developing and the blastula contains an inverted organization of cells in terms of the extent of their cleavage stage development. This allows us to assess how blastula patterning affects development without concerns about residual effects from the chamber. In short, the temperature gradient is a transient event and does not embed a division history or memory in the cells. We conclude it is feasible to repattern blastula tissue organization – cell size gradient – via spatial control of the cell division period. Thus, we next sought to determine the effect of blastula cell size gradient inversion on spatiotemporal patterning of ZGA.

### Inverting the blastula cell size gradient inverts the patterned onset of zygotic genome activation

ZGA onset in cells of vertebrate blastula embryos is coupled to achieving a threshold cell size or DC ratio; also called NC ratio [23–25, 27, 29]. In *Xenopus*, ZGA normally occurs first in AP cells because they divide faster and reach the cell size threshold faster than VP cells (**Fig. S2A**) [23]. We therefore sought to test the hypothesis that blastula cell size organization determines the patterning of ZGA, utilizing our inverted embryos. We utilized 5-ethynyl uridine (EU) metabolic labeling to image nascent transcription in blastomeres of whole-mount embryos, as was done previously [23]. The inverted embryos show a clear spatiotemporal inversion of the onset of large-scale ZGA (**Fig. 2A**). Nascent transcripts are detectable first in small cells of the VP of the inverted embryos, while the AP cells remain inactive, at the late blastula stage. In contrast, control embryos show the normal AP-first ZGA pattern. By imaging the inverted embryos as they further develop, we find that the temporal ordering of ZGA is flipped. It occurs first in VP, next in equatorial regions, and finally in the AP cells (**Fig. 2B**). Similarly, if we apply gradients along a horizontal axis, we can create horizontal gradients of cell size and ZGA onset, such as along the equatorial region (**Fig. S2B-E**).

**Figure 2.**
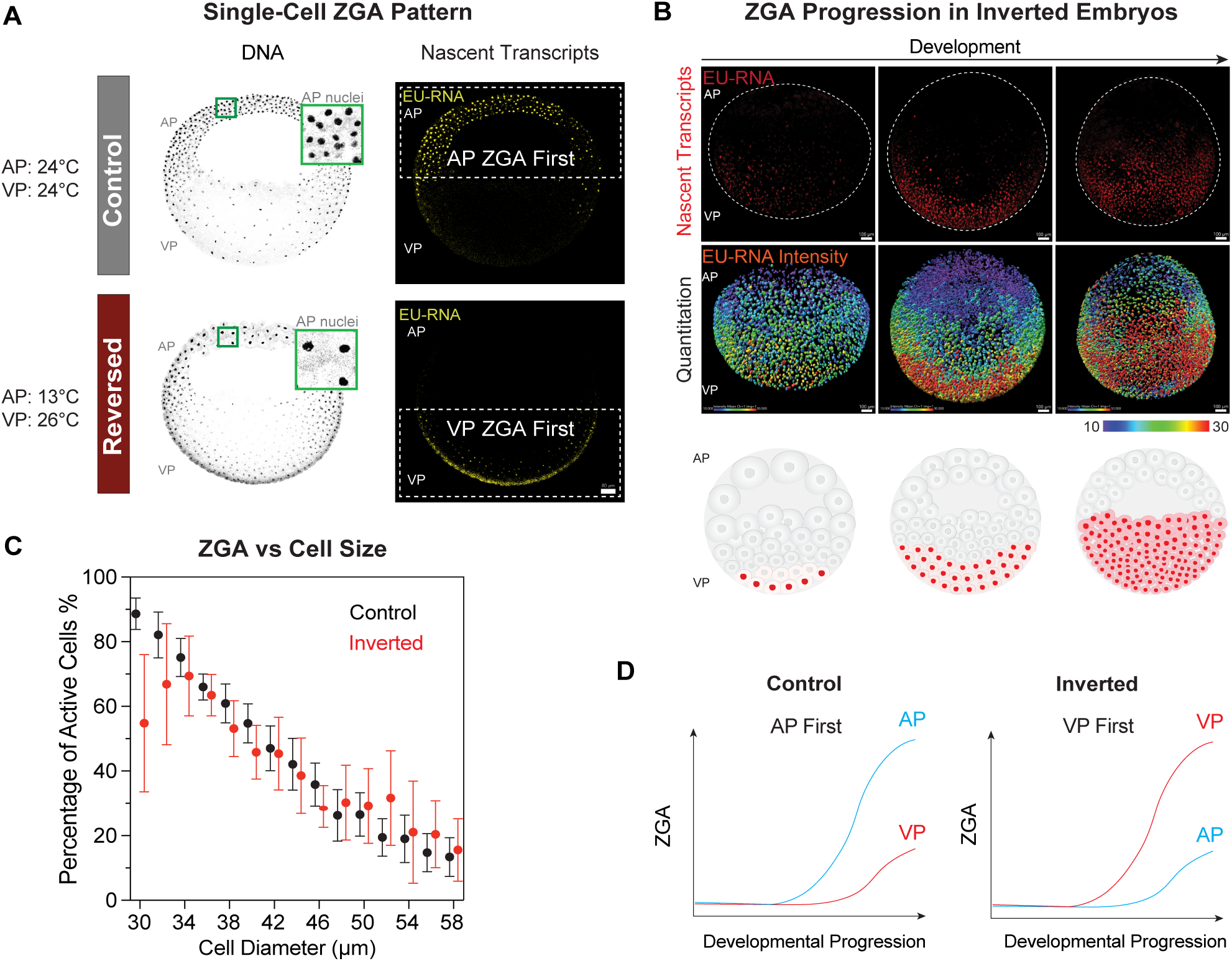
Inversion of the blastula cell size gradient flips the spatiotemporal ordering of genome activation. **(A)** Confocal slices from image stacks of whole-mount embryos at 10 hpf, showing DNA (inverted grayscale) and nascent transcripts (EU-RNA, yellow). Box shows enlarge view of nuclei and inter nuclear distance (cell diameter). Cell size gradient and pattern of ZGA are inverted in 13°C/26°C embryo at late-blastula stage. Min-max value adjusted to the same in DNA and EU-RNA labeling. Scale bar, 80 µm. **(B)** Bottom-up onset of ZGA within inverted embryos fixed at 11 to 12 hpf. In contrast to control embryos which show top-down ZGA [23], within inverted embryos nascent transcription initiates on VP and then progresses over time toward AP cells. Fluorescence confocal images of nascent transcripts (EU-RNA, top) and heatmap showing quantitation of EU-RNA per nucleus (bottom): low transcription (gray) to high (red). Dashed lines show the boundary of embryos. Scale bar, 100 µm. **(C)** Percentage of transcriptionally active cells in late blastula embryo as a function of cell diameter. Mean value within 2 µm bins of cell diameter. Mean +- SEM, N=4 embryos for 23°C, N=3 embryos for 13°C/26°C. **(D)** Scheme showing spatial and temporal onset of ZGA in control versus A-V inverted embryos. Control: nascent transcription detectable in AP cells prior to VP cells. Inverted blastula: nascent transcription detectable in VP cells prior to AP cells.

To determine whether these patterns of ZGA onset are governed by a cell size-dependent trigger, we quantified nascent transcription as a function of blastomere size. In both inverted and control embryos, when cells get smaller, they show an increase trend in nascent transcription (**Fig. 2C**), similar to what has been reported previously [23]. Additionally, the ZGA onset pattern in a side-to-side gradient embryo can also be explained by cell size-dependent onset (**Fig. S2F**). These data suggest that blastula cell size organization dictates ZGA spatiotemporal patterning, and this depends on a cell sizer mechanism, as has been shown for control embryos. Because the A-V inverted embryos show an inversion of the spatial onset of ZGA (**Fig. 2A, C, D**), we were positioned to directly characterize the contribution of ZGA patterning to development.

### Development requires proper temporal ordering of ZGA

Previous efforts to dysregulate cell division along developmental axes of early blastula or blastoderm were ultimately tolerated [28, 31–33]. We wondered whether the inversion of the spatiotemporal ordering of ZGA in the *Xenopus* blastula would be sufficient to alter embryo development. To assess whether animal-first transcription is required for normal development, we imaged embryogenesis over the first-day post-fertilization. Although the inverted embryos resumed the normal cell divisions (**Fig. S1G** and **Video 1**) – following the removal from temperature gradient, we found that by 1 day post-fertilization, a large portion of the inverted embryos had died, with dissociated white cells covering the surface of embryos (**Fig. 3A, B, S3A** and **Video 2**). Embryos grown in chambers at uniform 24°C developed normally, as did those in chambers at 13°C, although their timing was delayed, and showed nearly no embryo death at the neurula stage (**Fig. 3B**, **S3A**) [36]. Importantly, only the application of the temperature gradient to delay AP caused embryo death. Imposing a temperature gradient along a horizontal axis which creates a left-right ZGA onset gradient (**Fig. S2E**), or applying the temperature gradient in reverse along AV axes, causes no embryo death (**Fig. 3C, S1H**). These data demonstrate that neither the chamber device nor the temperature gradient per se has a negative influence on development. Instead, it is specifically the inversion of ZGA onset along the A-V axis that causes embryo death. Because death ratios varied between clutches of embryos, we next chose to monitor the development of individual embryos, so as to reveal its correlation to the extent of slowdown of AP cell division and ZGA onset. In short, we sought to determine how the delay of AP ZGA relative to VP could explain embryo outcomes. We extrapolated the cell division number in the animal pole based on the surface cell density and calculated the relative delay in AP divisions compared to VP. We assumed the vegetal side of all embryos had similar cell cycle progression based on our imaging data (**Fig. 1D, E, S1H**), and the fact that the temperature of the plate proximal to VP was kept at 24°C both for temperature gradient and control chambers. We then plotted the division delay relative to embryo death and fit the data with logistic regression (**Fig. 3D**). We found that embryo death is dose-dependent, set by the extent of delay between AP and VP. When the AP delay exceeds 1.9 hours or ∼ 4 cell divisions, embryos are more likely to die than survive. These data show that proper patterning of ZGA onset is essential for embryo development. Although embryos can tolerate spatial delay or mismatch of ZGA along an equatorial axis (**Fig. S2D, E**), or further delay of ZGA in VP (**Fig. 3C**), they must undergo AP-first zygotic transcription to develop normally.

**Figure 3.**
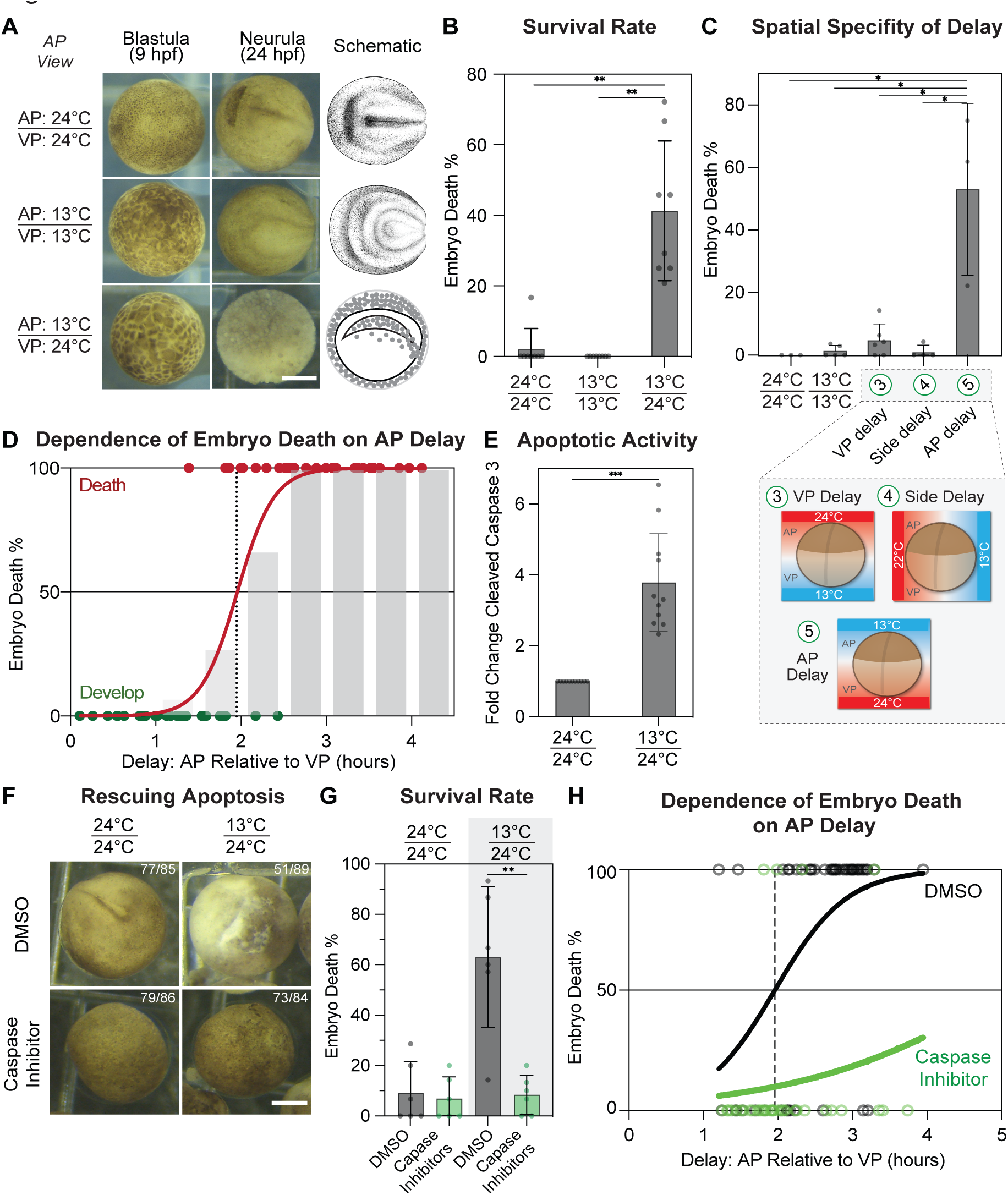
Altered ZGA patterning induces embryo death via apoptosis dependent on the extent of AP delay **(A)** Representative stereomicroscope images of developing embryos at 9 hpf and 24 hpf. 24°C/24°C control embryos are normal at 24 hpf (neurula stage) and 13/13°C embryos are normal but delayed. 13°C/24°C embryos are dead at 24 hpf, covered by white cells. *Xenopus* illustrations© Natalya Zahn (2022). Scale bar, 0.5 mm. **(B)** Quantitation of percentage of embryo death at 24 hpf following blastula incubation in chambers at indicated temperatures. N=8 experiments for each different temperature. Mean +- SD. N=12 or 24 embryos per experiment. Multiple paired parametric t-tests, P=0.0003 or 0.0006. **(C)** Embryo death at 24 hpf depends on direction in which temperature gradient is applied. (3) VP delay: 24°C/13°C (AP/VP), (4) 13°C|22°C (left/right), (5) 13°C/24°C (AP/VP). Percentages of embryo deaths. Only AP delay induces potent embryo death. Mean +- SD. N=3-6 experiments for each treatment. N=25-40 embryos per experiment. Multiple unpaired parametric t-tests, P=0.03, 0.004, 0.003, and 0.01. **(D)** Dose-dependence of embryo death dependent on extent of AP delay relative to VP. Scatter plot: embryos either alive or dead at 24 hpf as a function of AP division delay. Each dot represents one embryo. Data fit with logistic regression curve (red line). Overlay: Average percentage of embryo death per 0.5 hr delay bin (gray bar). Likelihood ratio test, ****, P<0.0001, significant. **(E)** Levels of cleaved caspase 3 relative to β-tubulin for embryos (26 hpf) quantified from Western blot images. Shown as fold change relative to 24°C chamber control embryos. Each dot represents one Western blot. Samples collected from two independent experiments. N=10 embryos pooled in each sample. Mean +- SD. Ratio paired parametric t-test, P<0.001. **(F)** Caspase inhibitors effectively rescue the apoptosis in inverted embryos (13°C/24°C temperature gradient) at 24 hpf. Scale bar, 0.5 mm. **(G)** Quantitation of percentage embryo death upon injection of caspase inhibitor. Mean + SD. N=6 experiments for each treatment. N=9-16 embryos per experiment, with an average of N=14.3 embryos. Paired parametric t-test, P=0.0024. **(H)** Dose-dependence of embryo death dependent on AP delay is desensitized by caspase inhibition. Scatter plot embryos either alive or dead at 24 hpf. Likelihood ratio test, ***, P=0.0009 for 13°C/24°C, significant; P=0.3149 for adding inhibitors, not significant.

To determine the nature of embryo death, we investigated whether delayed transcription in the animal pole induces embryo apoptosis. Previously, it was shown that transcription inhibition in *Xenopus* embryos leads to apoptosis and embryo death in the early gastrula [37]. We used cleaved caspase-3 as a marker of active apoptosis and found that many cells of the inverted embryos showed positive staining at 1 dpf (**Fig. S3B**). Those dead cells either accumulate inside the blastocoel or break into perivitelline space, which is why the embryo surface is covered by floating white cells in stereomicroscopy (**Fig. 3A, S3B**). In control embryos, no active caspase-3 staining was observed. Additionally, western blots confirmed a higher level of cleaved caspase 3 in the inverted embryos compared to control embryos (**Fig. 3E, S3C**). We reasoned that if embryos were actively dying via apoptosis from the AP ZGA delay, we could block it using apoptosis inhibitors. Indeed, injection of a cocktail of caspase inhibitors into 1-cell stage embryos before the temperature gradient treatment reduced apoptosis and resulted in significantly lower death ratios (**Fig. 3F, G**). Furthermore, when we plotted development outcome versus relative delay between AP and VP, we found that caspase inhibitors effectively rescued embryos from death, even when AP ZGA was largely delayed (**Fig. 3H**). The embryos do not develop further but are spared from an active cell death process within the first 1 dpf.

Together, these results suggest that embryos require a stereotyped spatial ZGA pattern for normal development. Delaying ZGA in AP relative to VP induces embryo apoptosis in a dose-dependent manner and can be rescued by caspase inhibitors. This process is specific for AP- delay and embryos are robust to all other perturbations to ZGA patterning. We found this compelling and quite distinct from the results of previous studies in which embryos tolerated cell division delays [28, 31–33] and suggested proper temporal ordering of ZGA is necessary to pattern the early embryo.

### Embryo quality control response is triggered by subregional delays in ZGA

Apoptosis is inactive in the blastula and embryos do not pause cleavage stage divisions in response to DNA damage [10]. Instead, in the early gastrula, the embryo triggers apoptosis following accumulated DNA damage or absence of embryo-wide transcription as part of a quality control response, also termed early gastrula checkpoint [37] (**Fig. 4A**). The theory is that development requires a zygotic anti-apoptotic program to counteract or balance the maternal pro-apoptotic protein. If not opposed by the zygotic program, apoptotic factors and activities would accumulate to sufficient levels to trigger apoptosis in early gastrula. We confirmed that the global transcription block causes embryo death via apoptosis (**Fig. S4A-D**). Next, we wondered whether the embryo could also sense spatially dysregulated transcription, such as loss of transcription in AP cells. We also reasoned that it would be useful to test a second method for delaying AP ZGA transcription and assess its effect on development.

**Figure 4.**
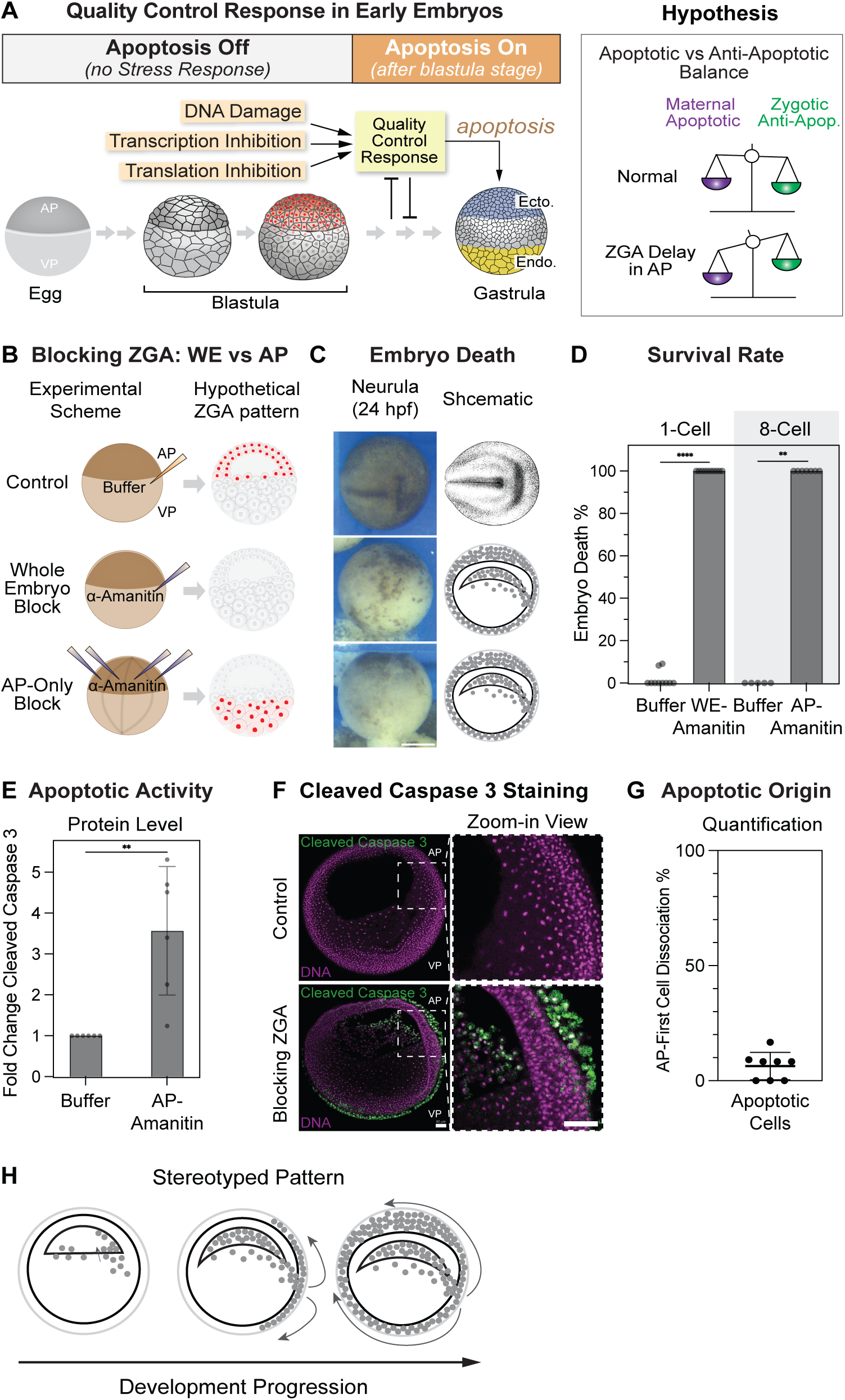
Embryo quality control responds to regionalized AP ZGA inhibition and initiatives apoptosis outside of AP **(A)** Schematic (top) of quality control response which senses stress (DNA damage, transcriptional inhibition) and triggers apoptosis; active after blastula stage. Schematic (bottom): temporal accumulation of maternal pro-apoptotic activity requires balancing by zygotic gene expression products. In embryos with AP ZGA delay the imbalance tips toward apoptosis. **(B)** Schematic representation of testing whole-embryo vs. regional ZGA delay. From α-amanitin or TBS control injection into 1-cell embryo (whole embryo RNAPII inhibition) or four AP cells at the 8-cell stage (regionalized AP RNAPII inhibition), with the hypothesized spatial patterns of ZGA. **(C)** Representative stereomicroscope images and schemes showing development outcome at 24 hpf (neurula stage). Drawing of the neurula: *Xenopus* illustrations © Natalya Zahn (2022). **(D)** Percentage of embryo death for each group, without or without global or regionalized transcriptional inhibition at 24 hpf. Mean +- SD. N=5-8 experiments. N=6-36 embryos per experiment. Mann Whitney test, P<0.0001 or P=0.0013. **(E)** Levels of cleaved caspase 3 relative to β-tubulin from Western blot quantitation, shown as fold change relative to TBS-injected embryos (20 hpf). Each dot represents one Western blot. Samples collected from three independent experiments. N=10 embryos per sample. Mean +- SD. Ratio paired parametric t-test, P=0.004. **(F)** Confocal slices from image stack of whole-mount embryos showing position of apoptotic cells. 14 hpf (gastrula stage). DNA staining (magenta) and cleaved caspase 3 staining (green). α- amanitin-injected embryos show apoptotic cells filling blastocoel and in perivitelline space but largely absent from AP tissue. Scale bars, 80 µm. **(G)** Quantitation of stereomicroscope movies showing location from which apoptotic cells are initially released from dying embryos. Percentages of embryos with dissociated cells first seen from the AP region. Each dot represents percentage in one experiment. Mean +- SD, N=8 experiments, N=10-12 embryos in each experiment. **(H)** Schematics summary of the stereotyped progression of apoptosis inferred from live imaging and from confocal imaging of whole-mount temperature gradient-treated and α-amanitin-injected embryos.

To test whether the subregional delay of ZGA in AP cells can be sensed by the embryo quality control response, we sought to specifically block transcription just in AP cells. To do so, we injected polymerase II inhibitor α-amanitin directly into the 4 animal blastomeres in 8-cell stage embryos (**Fig. 4B**). This would allow us to evaluate developmental consequences of regionalized transcription inhibition, compared to global transcriptional inhibition. We found that when control embryos developed to the neurula stage at 24 hours post-fertilization (hpf), embryos whose AP cells were transcription inhibited were mostly dead, similar to those with whole-embryo transcriptional inhibition (**Fig. 4C, D**). This result indicates that the embryo quality control response can indeed sense alternations to the spatiotemporal onset of ZGA, and in particular a block to ZGA in cells in the animal cap.

To determine the nature of embryo death following AP transcription block, we performed immunostaining of cleaved caspase-3 in transcription-inhibited embryos. Similar to the inverted embryos from temperature gradient treatment, these embryos exhibited apoptotic cells inside blastocoel as well as dead cells breaking into perivitelline space outside the embryo, trapped by the transparent vitelline membrane (**Fig. 4F**), as well as elevated levels of apoptosis via western blot (**Fig. 4E**). To our surprise, the animal cap cells appeared normal, not apoptotic, even though they were transcriptionally blocked. This suggests quality control response may be non-autonomous as the embryo can sense improper ZGA in AP cells and trigger death elsewhere.

To determine the stereotyped process of embryo death actively triggered by embryo quality control response, we analyzed time-lapse images. We found that white or dead cells initially appeared on the sides or bottom of the embryos, rather than directly from the animal cap (**Fig. 4G, H, S4E**, and **Video 2**). Dead cells may be released from equatorial or vegetal regions. Moreover, the structure of animal caps remains organized in embryos in which apoptotic cells accumulate in the blastocoel and perivitelline space, suggesting that apoptosis may not originate in the animal cap. To further understand the origin of apoptotic cells, we lineage traced the progeny of specific embryonic regions labeled at the 32-cell stage. We labeled B1 or D1 cells on the dorsal side of the embryo with dextran-488 (**Fig. S4F**) [38]. Progeny of B1 migrates downward and becomes the equatorial region on the dorsal side, while descendants of D1 become the center and bottom of the vegetal mass. We hypothesized apoptotic cells should originate from either the equatorial regions or vegetal regions or both. We confirmed proper regionalized labeling in blastula at 5.5 hpf via fluorescence microscopy. From top views, the daughter cells of B1 formed a patch on the side of the animal pole (**Fig. S4G**) as expected. For D1-labeled embryos, bottom views showed a patch of green cells on the edge of the vegetal pole (**Fig. S4H**), as expected [38]. To determine whether apoptotic cells originate from B1 or D1, we imaged transcription-inhibited embryos at 12 hpf and measured whether lineage-traced cells comprised the apoptotic pool in the blastocoel or perivitelline space. For ZGA-blocked embryos, only B1-labeled embryos showed fluorescence dissociated cells in the peri-vitelline space (**Fig. S4G, I**). Although D1-labeled ZGA- blocked embryos also showed massive cell dissociation, these cells were not fluorescent (**Fig. S4H, I**). Furthermore, because dissociated cells also accumulate in the blastocoel, we cut open the embryos to examine this population of dead, dissociated cells. We found many fluorescent cells only in B1-labeled α-amanitin-injected embryos, but not in D1-labeled embryos or the embryos with normal transcription (**Fig. S4J, K**). As established earlier, dissociated cells are apoptotic (**Fig. 3E**). In total, 80% of ZGA-blocked embryos showed apoptotic cells originating from B1, while only 15% showed apoptotic cells originating from D1 (**Fig. S4I**). Moreover, we tracked the descendants of C1 and C4 cells, which also populate later equatorial regions, and found that 80% and 50% of embryos showed fluorescent apoptotic cells (**Fig. S4L-Q**). These results suggest that apoptotic cells induced by transcription inhibition primarily originate from the equatorial regions rather than the vegetal mass.

### Nature of non-autonomous signal governing embryo quality control and development

In the early to middle blastula stage, apoptosis is inhibited by maternal proteins, such as Bcl2. After zygotic transcription begins, zygotic protein takes over the inhibitory role [39]. To determine the nature of the quality control response that actively kills the embryo (but not necessarily ZGA-delayed cells), and whether it is pro- or anti-apoptotic we sought to use cultured explants. We hypothesized that the presence of a signal from AP to the equatorial region in embryos would either promote or inhibit apoptosis and embryo death (**Fig. 5A**). Explants of *Xenopus* animal cap and vegetal mass from the late blastula are capable of continuing to undergo developmental events [10, 11]. They have been used to characterize the nature of secreted signals such as Wnt signaling and to differentiate between autonomous and non-autonomous phenotypes/responses [40, 41].

**Figure 5.**
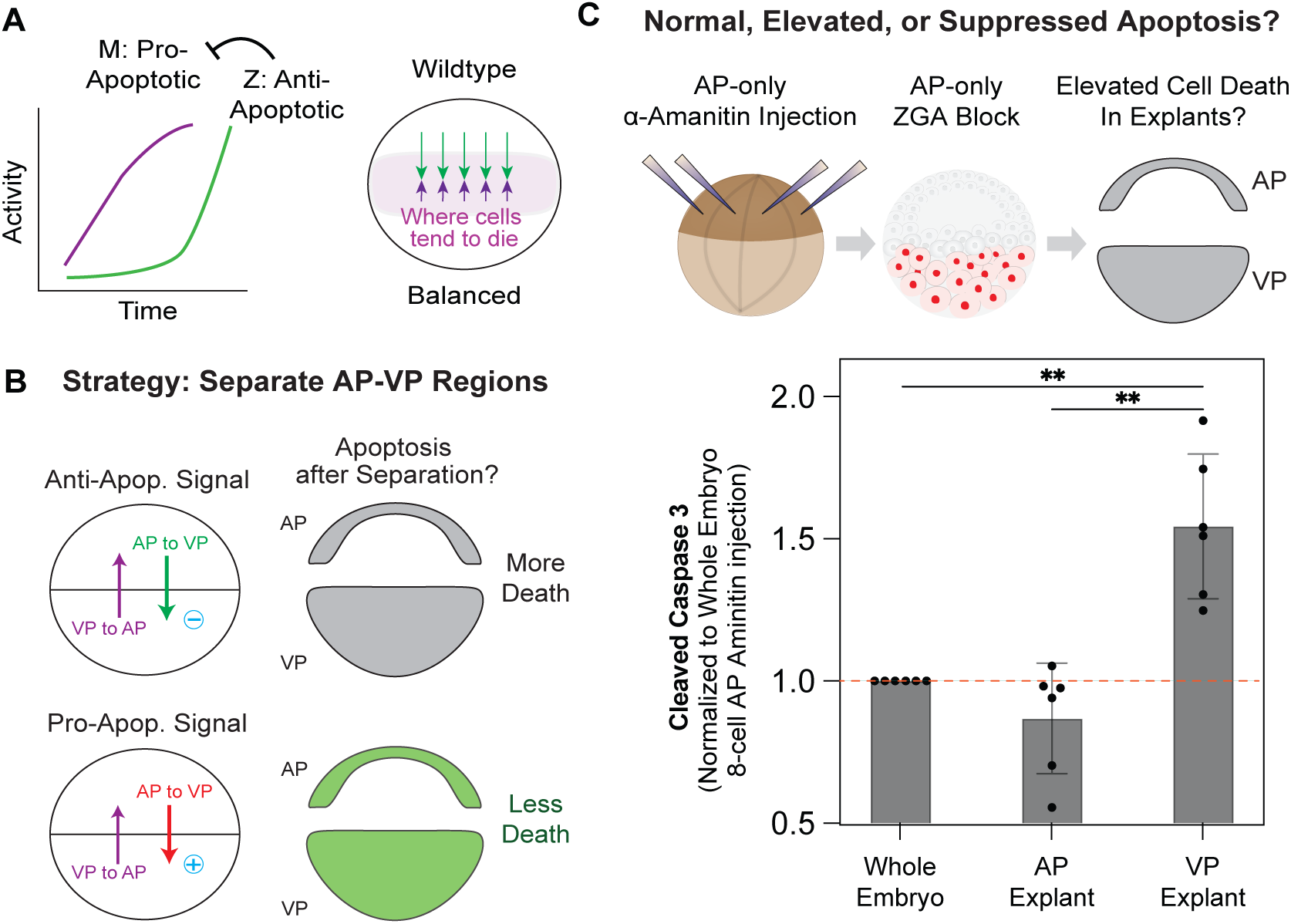
A non-cell autonomous signal underlies embryo quality control. **(A)** Schematic representation of zygotic repression of apoptosis and hypothetic spatial partitioning of signals for embryo quality control following ZGA. **(B)** Potential hypotheses: pro- or anti-apoptotic signals between animal pole and vegetal mass, along with predictions of potential developmental outcomes once explanted tissues are separated from each other. **(C)** Top: experimental overview: microinjection of α-amanitin into four animal pole cells at the 8- cell stage, with the expected spatial pattern of ZGA. Bottom: Fold change of relative level of cleaved caspase 3 of Western blot in explants compared to whole embryos (20 hpf). All sample injected at 8 cell stage in AP blastomeres. Each dot represents one Western blot. N=10 embryos pooled in each sample. Ratio paired parametric t-test, P=0.001 and 0.002.

To test this hypothesis, we cultured explants in embryos in which AP transcription was blocked. We dissected the embryos into animal caps and vegetal bottoms at 7 hpf, stage 8 (**Fig. 5B**), and collected explants and embryos for analysis of apoptosis at 20 hpf. In embryos with transcription inhibition on the animal side, the vegetal explants showed significantly higher levels of apoptosis compared to the animal explants and whole embryos (**Fig. 5C**). These results suggest that a non-autonomous anti-apoptotic signal prevents apoptosis in equatorial regions (part of the vegetal hemisphere). It also explains how embryo death can originate in regions of embryos in which transcription was not blocked, while AP-ZGA-blocked cells do not appear dead. We surmise that embryo quality control can sense AP ZGA delays and actively initiate embryo death in equatorial regions to eliminate those with improper ZGA patterning.

## Discussion

ZGA onset is highly stereotyped within an organism, occurring at a reproducible time, upon achieving a DC or NC ratio, or following a set number of divisions [6–8, 13]. In embryonic models that contain a blastula cell size gradient, the onset of zygotic transcription follows a precise spatial and temporal order - cells initiate large-scale ZGA dependent on reaching a threshold cell size or DNA-to-cytoplasm ratio [23]. In this study, we reveal that the spatiotemporally patterned onset of ZGA along the animal-vegetal axis is absolutely essential for embryo development.

The early embryo is highly plastic and resistant to perturbations to its tissue organization. Removal of half of the embryo at the 2-cell stage or blastula stage in sea urchin, frog, and mouse embryos initiates compensatory growth and ultimately embryos undergo normal development [42–47]. Moreover, previous studies in fly, wasp, and frog have shown that the embryo tolerates and compensates for an alteration of cell division cycles to one side of embryonic tissue [28, 30–33, 46]. In our study, embryogenesis proceeds normally even when creating spatial delays in cell division on one side of the embryo or by further accelerating cell division in the animal pole while delaying it in vegetal pole. However, we discovered that the embryo is highly sensitive to delay the onset of ZGA in AP cells relative to the VP cells. An embryonic quality control response is triggered dependent on the extent of AP ZGA delay and this process can be recapitulated by inhibiting RNAPII-dependent transcription just in AP cells. Thus, although the embryo is highly malleable and robust, AP-first zygotic transcription is essential to development in this vertebrate model.

Embryo quality control responses have been incompletely described in *Xenopus*, zebrafish, and mouse [10, 12, 39, 48, 49]. Generally, in non-placental embryos, blastomeres divide rapidly in blastula stages without gap phases or checkpoints [1, 3, 4]. DNA damage from irradiation does not pause cell divisions as it would in somatic tissues [10]. Instead, the embryo evaluates the extent of damage or loss of quality in early gastrulation. Global block of transcription throughout the embryo triggers quality control and embryo apoptosis [10, 11]. We now further demonstrate that *Xenopus* embryos can sense and trigger quality control response from regionalized delays in ZGA. Investigating the nature of the quality control response we reveal that it is non-autonomous: transcriptionally inhibited or ZGA-delayed AP cells do not appear to die first and instead embryo death appears to originate from equatorial regions and involves release of apoptotic cells into blastocoel and perivitelline space. Explant experiments suggest the existence of anti-apoptosis signals from the animal pole that help the embryo to pass quality control.

Early embryos are quite robust systems to physical perturbations including mismatch in cell division speeds, tissue removal, and in this current study tolerant to even some spatiotemporal alterations of ZGA timing along certain embryonic axes. This robustness however ultimately revealed a key function of patterned ZGA onset (AP-first), and that the embryo has a quality control system to sense it. This quality control mechanism enables the embryo to monitor the progression of genome activation and abolish development once ZGA mispatterning is too extended to recover. We anticipate future studies to leverage the embedding of ectopic ZGA gradients on embryos from different species to reveal unappreciated roles for quality control in early development.

## Materials and Methods

### Experimental model and subject details

African claw frogs *Xenopus laevis* were used for egg collection. All animal experiments in this study were performed according to the Animal Use Protocol approved by the University of Pennsylvania Animal Care and Use Committee.

### Animal Husbandry and Ovulation

Mature *Xenopus laevis* females and males were purchased from Nasco or Xenopus 1, and they were maintained at 20°C in tanks of a recirculating aquatic system. During the experiment, no harm was caused to the animals. The females were used for acquiring eggs and males were used for preparing sperms, the procedures of which have been described previously [23, 50]. To prime female frogs for ovulation, 100 U of pregnant mare serum gonadotropin (PMSG) was injected into the dorsal lymph sac 4-7 days before the experiment. Then, female frogs were injected with 500 U of human chorionic gonadotropin (HCG) at 14-18 h before egg collection and kept at 16°C in 2 liters 1x Marc’s Modified Ringer’s (MMR: 100 mM HEPES pH 7.8, 2 mM EDTA, 2 M NaCl, 40 mM KCl, 20 mM MgCl2, and 40 mM CaCl2 in dark. After egg procurement, the females were quarantined in high marine salt for at least four days before returning to the recirculating aquatic system. The ovulated females were not used until rest for at least 3 months. To prepare testes, male frogs were euthanized with 0.2% tricaine in 2 liters deionized water for at least 20 min before dissection. The testes were kept in L-15 medium on ice for same-day use or stored in a 4°C room for up to 1 week.

### *In vitro* fertilization (IVF)

All IVF in this study were performed at 16°C or 23°C. The procedures for IVF have been described previously [23, 50]. Briefly, female frogs were gently squeezed to obtain the eggs in a glass dish.

0.6 ml of sperm slurry was evenly added to a monolayer of eggs. The sperms and eggs were thoroughly mixed by gently sliding a plastic pestle. Five minutes after adding sperms, the dish was flooded with 15 ml of 0.1 x MMR to facilitate fertilization. At 20 minutes post-fertilization (mpf), fertilized eggs turn the pigmented animal side to the top. At 30 mpf, the embryos were incubated with 3.3% L-cysteine in 0.1xMMR for 3-5 min, with gentle swirling of dish to remove the jelly coats. Three rounds of 0.1xMMR wash were performed to remove residual L-cysteine, and embryos were kept in 0.1 X MMR for further use.

### Microinjection

The procedures for microinjection have been described previously [23, 50]. Briefly, embryos at the 1-cell stage (40-50 mpf) were transferred into a plastic dish containing 3% Ficoll in 0.5 x MMR. For 5-ethynyl uridine (EU), embryos were microinjected with 10 nl of 50 mM EU using a PLI-100 picoliter microinjector (Medical Systems, Greenvale, NY). The final concentration of EU inside embryos was ∼ 0.5 mM.

For α-amanitin, 10 nl of 0.1 or 2 ng/μl solution was injected into 1-cell embryos for a final concentration of 1 or 20 pg/μl. For animal pole injections, 2 nl of 0.2 ng/μl to 1 ng/μl was injected into each 4 animal pole cells, for a final concentration of 16 pg/μl to 80 pg/μl. All the concentration induced 100% embryo death.

For caspase inhibitors, a cocktail of 6 inhibitors was injected into 1-cell-stage embryos. The final concentration was 80 μM of Ac-DEVD-CHO, 16 μM of Z-DEVD-FMK, 4 μM of Z-VAD-FMK, 13.3 μM of GSK-872, 6.7 μM Nec-1, and 6.7 μM VX-765.

In lineage tracing of B1, C1, or C4 cells, 2 nl of 1 μg/μl Dextran Alexa-488 dye was injected into each of the 2 cells at the 32-cell stage, for a final concentration of 110 ng/μl.

In lineage tracing of D1 cells, 2 nl of 4 μg/μl Dextran Alexa-488 dye was injected into each of the 2 D1 cells at the 32-cell stage, for a final concentration of 110 ng/μl.

### Generating AP-to-VP temperature gradient

Physical Set Up. The temperature gradient device includes a chamber, temperature-controlling machinery, and aluminum heat sinks [28]. The chamber was assembled by a U-shaped acrylic insert sandwiched between two aluminum blocks. Each block contains a hole on the side, on which a thermistor was inserted for temperature measurement, sealed by electrical tape. 40 embryos were transferred to the top of the bottom block with an acrylic insert holding the 0.1x MMR. The top block was gently added to the top of the embryos. 0.1x MMR was added on the open side of the U-shaped acrylic insert to reduce the air bubble to a minimum. A parafilm stripe was used to wrap the chamber to provide mechanical strength for assembly and to minimize water evaporation. After wrapping two rounds of parafilm, the assembled chamber was moved to a plastic container with two shallow dishes filled with water on the side, to increase the moisture, wehich we found essential for maximum embryo survival. A Peltier cooler/heater was placed at the bottom and the top of the assembled chamber, directly affixed to each aluminum block , and to each Peltier a heatsink was added to facilitate heat dissipation. The thermistors and Peltier elements on each side were linked to a digital temperature controller separately, which allows precise and independent control of the temperature of each aluminum block. Before the temperature gradient, the embryos were incubated at 23°C and we sorted them for proper number of cleavages. Embryos were transferred into a 6-well plate containing 0.1x MMR. Each well of the 6-well plate contained 40 embryos for each temperature chamber. The AP-to-VP temperature gradient was started at the 8-cell stage once animal and vegetal cells have become physically separated, via the 3^rd^ embryo cleavage.

Temperature Control and Monitoring. After assembly of the temperature gradient device, dry ice pellets were added to the top of the top heatsink every 15 min to dissipate the heat generated by the Peltier element functioning to cool down the top block. For control chambers, we set temperature controllers to 24°C/24°C using all the same chamber elements. For control embryos incubated13°C/13°C, we placed the assembled chamber on a 12°C cold plate whose temperature is maintained by a circulating water chiller. The temperature of each block was also measured by the thermistor and shown to be stable. Each temperature controller was calibrated to the mean temperature of 0.1x MMR before first use. During treatment of temperature gradient or even temperature, the temperature of each block was recorded every 15 min to ensure a stable temperature during treatment. The temperature of the top block of 13°C/24°C chamber took 5-10 min to cool down to 13°C, while the bottom block was maintained at 24°C.

Chamber Disassembly and Setup for Imaging. The temperature gradient ended before 9 hpf (stage 10), to maximize the cell division delay in animal pole cells compared to vegetal cells, but also to ensure initial cell movement in gastrulation was not affected. Before removing the final piece of parafilm wrapping the open side of the insert, we looked from the side of the transparent acrylic insert to make sure the orientation of embryos was normal and there was no large air bubble inside. After the parafilm was removed, the chambers were opened with the top side facing down to a large glass dish containing 0.1x MMR. Gravity cause embryo transfer to the dish. We changed the 0.1x MMR to remove any debris from broken embryos. Embryos were imaged in groups and used for fixation or development tracking at 23°C.

### Generating side-to-side temperature gradient

Physical Set Up. The side-to-side temperature gradient uses the same temperature-controlling machinery and heatsinks, except the chamber was assembled by two elongated aluminum blocks with an acrylic insert in between. Each block contains a hole on the short side, on which a thermistor was inserted for temperature measurement, sealed by electrical tape. The chamber was assembled first, and another long piece of electrical tape was used to wrap the side and bottom of the chamber. The acrylic insert has an opening on the top, and 0.1x MMR was added to the opening. 20 to 30 embryos were added from the top to form a horizontal line with no embryos sitting on top of others. A piece of glass plate was added to cover the top of the opening to reduce evaporation, and 0.1x MMR was added every 30 min from the side to prevent drying of the reservoir. Peltiers were sandwiched between a large heatsink and the long side of aluminum blocks. The side-to-side temperature gradient started at the 8-cell stage.

Temperature Control and Monitoring. After the assembly of the whole device, dry ice pellets were replenished on the top of the heatsink of the cooling side every 15 min to provide maximum dissipation of heat generated by the Peltier. For control chambers, we ran 22°C | 22°C by placing the assembled chamber on a room temperature bench without the temperature controller machinery. The temperature of each block was also measured by the thermistor. Each temperature controller was calibrated to the mean temperature of 0.1x MMR before first use. During temperature gradient treatment, the temperature of each controller was recorded every 15 min to ensure stable temperature treatment. The temperature of the block on the cool side took 5-10 min to cool down to 13°C, while the control block was maintained at 22°C. Note: 13°C-to- 22°C difference was the maximum gradient we could achieve due to the size of the reservoir.

Chamber Disassembly and Setup for Imaging. The side temperature gradient ended at 8 hpf (stage 9), prior to gastrulation cell movements. After, the glass plate and tape were removed, the chambers were opened with the top side facing down to a large glass dish containing 0.1x MMR. Gravity cause embryo transfer to the dish. Embryos that came from the same chamber were transferred to a well of 6-well plate containing 0.1x MMR. Embryos were imaged in groups and used for fixation or further development tracking at 23°C.

### Stereomicroscope imaging

Brightfield stereomicroscopy was performed using a Leica MZ FLIII microscope (Plan APO 1.0x objective) and MC170 HD camera, with LAS X software. Magnifications from 0.8x to 4.0x were used for imaging. Exposure time was adjusted between 30 to 100 ms under different light conditions and magnifications, and images were not saturate. In time-lapse imaging, images were taken every 1 min to 10 min, lasting from 2h to 18h. Calibration of scale was done by imaging the hemocytometer under different magnifications.

Fluorescence stereomicroscopy was performed under an Olympics MVX100 microscope (MV PLAPO 1.0x objective) and OP71 camera, with DP Controller software. Magnifications from 0.63x to 4.0x were used for imaging. Focus planes for imaging were adjusted under brightfield and not changed for fluorescence imaging. ISO was set to 400 and image intensity was an average of two images taken. Exposure time in brightfield was adjusted between 33 to 100 ms to not saturate. Exposure time for fluorescence imaging was set to 500 ms and images were mostly not saturated. Calibration of scale was done by imaging the hemocytometer under different magnifications.

### Cell cycle analysis of surface cells on the animal pole

We first determined the cell density in control embryos after each cell division. Then we inferred the cell cycle of embryos after temperature treatment based on their cell intensity.

First, we took a time-lapse of the top views of *Xenopus laevis* embryos to determine the timing of the 3^rd^ to 14^th^ cell division (average of 8 cells in each of N=3 embryos, unless for the 3^rd^ and 4^th^ division that there are less than 8 cells visible from the top views). Note that for the 14^th^ division, the cell cycle becomes asynchronous. After 6 minutes of average division timing in each embryo, cell density was determined by counting the cells in a center circle. Then, we multiply the cell number and a size factor (ratios between areas of embryo and center circle) to get the inferred total cell number in AP surface, considering embryos with large size should have larger cells after the same number of divisions. Note that it is not the real total number, as the embryo surface in the periphery of top views is tilted and may include more cells even with the same cell density.

Then we used log2(“inferred total cell number in AP surface”) in X axis and cell cycle number in Y axis to establish linear regression with a formula of, y = 1.5269x - 1.4366 and R² = 0.9924, indicating a high-confident linear relationship.

Note that although the cell number doubles after each cell division, some daughter cells are underneath others, making those cells not visible from the AP surface, so it takes around 1.5 cell cycles to double the cell number seen from the AP surface.

For the embryos after temperature chamber treatment, we counted and inferred the “total cell number in AP surface” and then used the formula to infer the cell cycle of AP surface cells in those embryos.

### Whole-mount click chemistry and immunofluorescence

Metabolic labeling of nascent transcripts and immunostaining have been described in detail previously [23, 50].

4% PFA/1 x MEM solution is made fresh and aliquoted into 2ml glass vials before fixation. At desired stages, embryos from the same treatment groups were transferred together into a 2ml glass vial (N=10-20 embryos each) using a transfer pipette. Glass vials were put to a rotator for low-speed rotations for 2h at 23°C or overnight (>12h) at 4°C.

After fixation, fixative solution was removed, and methanol was added for dehydration on a rotator. After 5 min, methanol was replaced, and this process was repeated twice. Glass vials were stored in methanol at -20°C fridge before processing.

Embryos were rehydrated by washing sequentially with 75%, 50%, and 25% methanol in 0.5x SSC (75 mM NaCl and 7.5 mM sodium citrate) for 5 min each, followed by washing with 0.5x SSC two times. Embryos were bleached in 0.5x SSC containing 5% formamide and 2% H2O2 for 2-4 h under light at 4°C room. Embryos were washed with 0.5x SSC for 5 min for two times, followed by washing with 1x TBST for 30 min for eight times. Embryos were washed two times with 1xTBS for 10 min each.

Embryos were incubated with 25 μM TAMRA-azide, 100 mM Tris-HCl pH 8.5, 1 mM CuSO4, and 100 mM ascorbic acid for 5-12 h at 23°C. Embryos were extensively washed with 1x TBST all day at 23°C by changing buffer every 1h, overnight at 4°C, and all day at 23°C by changing buffer every 1h. For blocking, embryos were blocked with 10% goat serum in 1xTBST for 1h at 23°C. Then embryos were incubated with TO-PRO-3 (1:300 dilution in 1xTBST) overnight at 4°C. Embryos were washed with 1x TBST all day at 23°C by changing the buffer every 1h. Embryos were completely dehydrated with anhydrous methanol by replacing it more than four times. Embryos were cleared in BABB (mixture of 1 part benzyl alcohol and 2 parts benzyl benzoate) for 12-24 h before confocal imaging.

For immunostaining of cleaved caspase 3, embryos were not injected with EU and most of the process was the same except skipping the click chemistry reaction and the washing afterward. After blocking, embryos were incubated with primary antibody against cleaved caspase 3 (1:400 dilution in 1xTBST) overnight. Embryos were washed with 1x TBST all day at 23°C by changing the buffer every 1h. Then embryos were incubated with TO-PRO-3 (1:300 dilution) and 2^nd^ antibody (1:200 dilution in 1xTBST) overnight at 4°C. Embryos were washed with 1x TBST all day at 23°C by changing the buffer every 1h. Embryos were dehydrated and cleared the same way as mentioned in the previous paragraph.

### Confocal microscopy

Confocal microscopy was performed with the ZEN software on a Zeiss LSM710 confocal microscope. EU-RNA, TO-PRO-3, and Cleaved Caspase 3 signals in *Xenopus* embryos were imaged with a frame size of 1,024 pixels x 1,024 pixels using lasers 561 nm (0.15% power), 633 nm (2% or 5% power) and 488 nm (5% power), respectively, without saturating signals. Whole embryo Z-stacks were collected using the Plan-Apochromat 10x / 0.45 objective with 5 μm intervals.

### Confocal image analysis for cell sizes and nascent transcriptions

Metabolic labeling of nascent transcripts and immunostaining have been described in detail previously [23, 50].

Briefly, the nuclei surface was rendered based on TO-PRO staining in Imaris 10.2 (Oxford Instrument). log2 of total nuclei number was used for staging the developmental progression of embryos. The nuclear volume was obtained directly from the statistics generated by the software. The 3D positional information, i.e., X, Y, and Z positions, were exported for calculating the vector distances between one nucleus and its three closest nuclei, and the average vector distance was used as a proxy of the cell diameter of that cell.

The original nuclear EU-RNA intensity was measured by overlaying the segmented nuclei on the EU-RNA-TAMRA channel. The cytoplasmic background was measured by generating a 1- 2 μm shell surrounding individual nucleus surfaces and overlaying the segmented nuclei on the EU-RNA-TAMRA channel. The net nuclear EU-RNA intensity was calculated by subtracting the cytoplasmic background from the original nuclear EU-RNA intensity, and the nuclear EU-RNA amount was calculated by multiplying the net nuclear EU-RNA intensity with the nuclear volume.

The Threshold of the mean EU-RNA intensity and amount at the 8000-cell stage (log2 cell number = 13) was used to calculate the percentage of transcriptionally active cells in individual embryos at various stages.

### Western blot and analysis

Embryos at stage 8 (6 hpf) were transferred to 1x MMR and dissected into animal caps and vegetal bottom based on pigment, with equatorial regions trimmed off. At stage 9 (10 hpf), N=10 explants or embryos were transferred into a 1.5 ml Eppendorf tube to contain all the cells that would be disassociated (animal caps with 60 μl 1xMMR, vegetal bottoms with 140 μl 1xMMR, no- vitelline embryos and normal intact embryos with 260 μl 0.1xMMR). At neurula stage (20 hpf), 2x sample buffer was added to each tube for sample collection, with an amount equal to the 0.1x or 1x MMR already in the tubes (2x sample buffer consists 47.5% of 2x lysis buffer, 47.5% of 4x SDS loading buffer and 5% 2-mercaptoethanol and protease inhibitor cocktail; 2x lysis buffer: 40 mM Tris-HCl pH8, 0.1 M NaCl, 4 mM EDTA, 0.2% NP-40; 4x SDS loading buffer: 0.2 M Tris-HCl pH 6.8, 40% glycerol, 8 % SDS and 0.02% bromophenol blue). The samples were pipetted 30 times with half the total liquid volume, using P200 or P1000 tips. The samples were heated for 2 min at 95°C and centrifuged at 13,000 rpm for 5 mins at 4°C. The supernatant of samples in the middle section of the tubes was aliquoted and kept at -20°C.

Before running the protein gels, equal amount of 4xSDS loading buffer was added to aliquoted samples to help with loading and running. 8 to 20 μl diluted samples were loaded onto each well of NuPAGE 4-12% 12-well Bis-Tris Gel. Bio-Rad Dual color standard was loaded to the first well of gel and 4x SDS buffer was loaded onto empty wells.

The gel runs at 85V for 20 minutes and 100V for 70 minutes at 23°C. Proteins on gels were transferred to a PVDF membrane (0.45 μm pore size, Millipore) with transfer buffer (10 mM CAPS and 20% methanol) at 20 V for 1 h at 23°C using a Bio-Rad Semi-Dry Transfer Cell. Membranes were blocked with 5% non-fat milk in 1 x TBST (20 mM Tris pH 7.6, 150 mM NaCl, and 0.1% Tween-20) for 1h at 23°C and incubated with rabbit anti-Cleaved Caspase 3 (1:500 dilution) primary antibody overnight at 4°C. Membranes were washed with 1xTBST 5 times for 5 mins each and incubated with secondary rabbit IRDye 680 (1:2500 dilution) for 1h at 23°C. Membranes were washed with 1xTBST 5 times for 5 mins each and rinsed with 1xTBS 5 times. Membranes were scanned under LI-COR Odyssey Infrared Imager (LI-COR Biosciences).

Because of the bleed-through of IRDye 800 secondary antibody in the 700 nm channel, membranes were first incubated with Cleaved Caspase 3 antibody (weak), stripped, and applied β-tubulin antibody (strong).

Membranes were washed 1 time for 2 minutes with 1xTBS and were incubated in stripping buffer for 15-30 minutes before rinsing with 1xTBS again 2 times for 3 minutes. Membranes were blocked with milk again for 1h at 23°C and incubated with mouse E7 anti-β-tubulin primary antibody (1:5000 dilutions) for 1h at 23°C. Membranes were washed with 1xTBST 5 times for 5 mins each and incubated with secondary mouse IRDye 800 (1:2500 dilution) for 1h at 23°C. Membranes were washed with 1xTBST 5 times for 5 mins each and rinsed with 1xTBS 5 times. Membranes were scanned under LI-COR Odyssey Infrared Imager (LI-COR Biosciences).

The signals of bands were analyzed using the gel analysis tool in Fiji (ImageJ2). The same size of boxes was drawn to include the whole lanes of β-tubulin and cleaved caspase 3. Plots of the amount of intensity were generated in Fiji and baselines were drawn underneath each peak of signals, accounting for different backgrounds around each band. The area of peaks was measured using the wand tool in Fiji and relative levels of cleaved caspase 3 were calculated by normalizing to the level of β-tubulin in each sample. In the figures, the relative levels were normalized to the levels in the control sample, and fold changes of the cleaved caspase 3 level were shown.

### Quantification and statistical analysis

All statistical parameters, including sample numbers, mean and standard deviation or error, were included in Figures and Figure legends. *p < 0.0332; **p < 0.0021; ***p < 0.0002; ****p < 0.0001; ns, not significant.

### Online supplemental material

Fig. S1 shows the effective temperature inside embryos during temperature gradient treatment and resuming of cell divisions after the treatment. Fig. S2 shows application of a horizontal temperature gradient induces an ectopic cell size gradient. This new gradient changes the genome activation pattern but not inducing embryo death. Fig. S3 shows transcription delay in animal pole induces apoptosis. Fig. S4 shows embryos can sense transcription delay in spatial subregions and that ZGA inhibition induces apoptosis in equatorial regions. Video 1 shows after transferring back to room temperature after temperature gradient treatment, the embryo resumes normal division rates (∼2 divisions/hour). Video 2 shows following temperature gradient mediated inversion, the embryo undergoes death at neurula stage.

## Data availability

All the data supporting this study are available from the corresponding authors upon request.

## Acknowledgements

We thank members of the Good, Klein, and Mullins labs at the University of Pennsylvania for helpful discussion and feedback. We acknowledge Dr. Lendert Gelens for the help with initial design of embryo temperature controller; Dr. Yuri Veklich for help on selecting the electronics; Peter Szczesniak and Jason Pastor of Penn MEAM Precision Machining Lab for fabricating aluminum devices and insert for embryo chambers, the Cell and Developmental Biology CDB Microscopy Core, RRID SCR_022373, for imaging support; Xenbase for useful resource on embryo images, staging, antibody, and gene expression; National *Xenopus* Resource (NXR) for training and guidance on frogs. This work was supported in part by National Institute of General Medical Sciences (R35GM128748), the March of Dimes, Charles E. Kaufman Foundation (M.C.G.)

## Author contributions

Conceptualization, W.Q., H.C., and M.C.G.; Methodology, W.Q., H.C., and M.C.G.; Resources, W.Q. and M.C.G.; Investigation: W.Q., H.C. and H.L.; Software, W.Q.; Formal Analysis: W.Q.; Writing – Original Draft, W.Q. and M.C.G.; Writing – Review & Editing, W.Q., H.C., H.L., and M.C.G.; Visualization, W.Q. and M.C.G.; Supervision, M.C.G.; Funding Acquisition, M.C.G.

Disclosures: The authors declare no competing interests.

---

**Table S1.** Key resource.

| REAGENT or RESOURCE | SOURCE | IDENTIFIER |
| --- | --- | --- |
| **Antibodies** |  |  |
| Cleaved Caspase-3 (Asp175) Antibody | Cell Signaling | Cat# 9661; RRID: AB_2341188 |
| Mouse monoclonal anti-tubulin | DSHB | Cat# E7-c; RRID:AB_528499 |
| Goat anti-Rabbit IgG (H+L) Secondary Antibody, Alexa Fluor 488 conjugate | Thermo Fisher Scientific | Cat# A11034; RRID: AB_2576217 |
| IRDye® 680RD Goat anti-Rabbit IgG Secondary Antibody | LI-COR | Cat# 926-68071; RRID: AB_10956166 |
| IRDye® 800CW Goat anti-Mouse IgG Secondary Antibody | LI-COR | Cat# 926-32210; RRID: AB_621842 |
| **Biological Samples** |  |  |
| *Xenopus laevis* embryos | This paper | N/A |
| **Chemicals, Peptides, and Recombinant Proteins** |  |  |
| Chorulon (HCG) | Midwest Veterinary Supply INC | Cat# 349.10450.3 |
| Pregnant Mare Serum Gonadotropin (PMSG) | BioVendor | cat# RP1782725000 |
| Ficoll® 400 | Sigma | Cat# F4375 |
| L-cysteine | Sigma | Cat# C7352 |
| Triton X-100 | Sigma | Cat# 10789704001 |
| Bovine serum albumin (BSA) | Sigma | Cat# A3059 |
| RNase-free water | Thermo Fisher Scientific | Cat# BP561-1 |
| Methanol | Thermo Fisher Scientific | Cat# A4524 |
| Methanol, anhydrous | Sigma | Cat# 322415 |
| Dimethyl sulfoxide (DMSO), anhydrous | Thermo Fisher Scientific | Cat# D2650 |
| 5-Ethynyl Uridine (5-EU) | Thermo Fisher Scientific | Cat# E10345 |
| Tetramethylrhodamine (TAMRA)-azide | Abcam | Cat# ab146486 |
| CuSO<sub>4</sub> | Sigma | Cat# 61230 |
| Ascorbic acid | Sigma | Cat# A7506 |
| TO-PRO-3 | Thermo Fisher Scientific | Cat# T3605 |
| Paraformaldehyde | EMS | Cat# 15710-S |
| Hydrogen peroxide | Sigma | Cat# H1009 |
| Formamide | Thermo Fisher Scientific | Cat# AC181090010 |
| Benzyl alcohol | Sigma | Cat# 305197 |
| Benzyl benzoate | Acros Organics | Cat# 105862500 |
| Ac-DEVD-CHO (N-Ac-Asp-Glu-Val-Asp-CHO) | Selleck Chemicals | Cat# S7901 |
| Z-DEVD-FMK | Selleck Chemicals | Cat# S7312 |
| Z-VAD-FMK | Fisher Scientific | Cat# PR-G7231 |
| GSK-872 | Selleck Chemicals | Cat# S8465 |
| Belnacasan (VX-765) | Selleck Chemicals | Cat# S2228 |
| Nec-1s (7-Cl-O-Nec1) | Selleck Chemicals | Cat# S8641 |
| Human PDGF-AA Recombinant Protein | Thermo Fisher Scientific | Cat# PHG0035 |
| NuPAGE™ Bis-Tris Mini Protein Gels, 4–12%, 1.0 mm | Thermo Fisher Scientific | Cat# NP0322BOX |
| Precision Plus Dual Color Protein Standard | Bio-Rad | Cat# #1610374 |
| Immobilon®-FL PVDF Membrane | Millipore | Cat# S1EJ084E03 |
| Instruments |  |  |
| PowerPac™ HC (High Current) | Bio-Rad | Cat# 1645052 |
| Trans-Blot® SD Semi-Dry Transfer Cell | Bio-Rad | Cat# 1703940 |
| LI-COR Odyssey 9120 Infrared Imager | LI-COR Biosciences | Cat# BZ2043001 |
| Experimental Models: Organisms/Strains |  |  |
| *Xenopus laevis* embryos | This paper | N/A |
| Software and algorithms |  |  |
| Fiji/ImageJ2 | NIH | RRID:SCR_002285,<br>https://fiji.sc |
| GraphPad Prism 10 | GraphPad | RRID:SCR_005375,<br>https://www.graphpad.com |
| Excel 2024 | Microsoft | https://www.microsoft.com/EN-US/microsoft-365 |
| RStudio | RStudio Inc | RRID: SCR_000432,<br>https://www.rstudio.com |
| Adobe Illustrator CC 2024 | Adobe | RRID: SCR_010279,<br>https://www.adobe.com/products/illustrator.html |
| LAS X | Leica | RRID: SCR_013673,<br>https://www.leica-microsystems.com/products/microscope-software/p/leica-las-x-ls/ |
| DP Controller | Olympus | https://www.olympus-lifescience.com/en/downloads/detail-iframe/?0[downloads][id]=847249636 |
| ZEN (black) | ZEISS | RRID: SCR_018163,<br>https://www.zeiss.com/microscopy/en/products/software/zeiss-zen.html |
| Imaris 10.2 | Oxford Instruments | RRID:SCR_007370,<br>https://imaris.oxinst.com |
| Xenbase, Zahn drawings | Xenbase | www.xenbase.org<br>RRID:SCR_003280 |
| Electric Elements for Temperature Gradient |  |  |
| Temperature Controller | Accuthermo | Model: FTC100D |
| H-bridge Amplifier | Accuthermo | Model: FTX100 |
| Thermistor | Accuthermo | Lot No: LSMN-TR2252-1M-WIRE |
| Peltier Thermo-Electric Cooling Module 6A | Jameco.com | Cat# 2265551 |
| RS-25-5 25W Single Output Switching Power Supply | Jameco.com | Cat# 323282 |
| RS-25-3.3 25W Single Output Switching Power Supply | Jameco.com | Cat# 693193 |
| 3-Conductor Power Cords with Pigtail (Unshielded) 290 Plug | Jameco.com | Cat# 38009 |
| 2-Conductor Line Cords with Pigtail | Jameco.com | Cat# 42042 |
| 22 AWG Black Solid Tinned-Copper Hook-Up Wire 25 Feet | Jameco.com | Cat# 9313-0-R-25 |
| 22 AWG Red Solid Tinned-Copper Hook-Up Wire 25 Feet | Jameco.com | Cat# 9313-2-25 |

**Table S2.** List of nonstandard abbreviations used in the paper.

| Phrases | Abbreviations |
| --- | --- |
| zygotic genome activation | ZGA |
| maternal-to-zygotic transition | MZT |
| animal pole | AP |
| vegetal pole | VP |
| DNA-to-cytoplasm ratio | DC ratio |
| nucleocytoplasmic ratio | NC ratio |
| animal-vegetal | A-V |
| 5-ethynyl uridine | EU |
| hours post-fertilization | hpf |
| day post-fertilization | dpf |
| diameter | dia |
| *in vitro* fertilization | IVF |

