## Supplementary Materials for "Spatially ordered zygotic genome activation fulfills embryo quality control"

**Online Supplemental Material**

Includes supplemental video legends and videos 1, 2 and supplemental figure legends, and Figures S1-S4

Video 1, related to Figure S1G

**Time lapse stereomicroscope imaging of cell divisions after application of temperature gradient and removal from chamber at late blastula stage**

A control embryo incubated at 24°C/24°C AP/VP (left) and embryo incubated in chamber with temperature gradient 13°C/24°C AP/VP (right). Imaged from 9:26 hpf to 15:26 hpf. View of animal pole surface. Images were collected every 4 minutes; display rate is 7.5 frames per sec (1 sec of video = 30 min real time). Scale bar 0.2 mm.

Video 2, related to Figure 3A, S4E

**Time lapse of cell dissociation and embryo death for blastulae whose cell organization was inverted in a temperature gradient device**

A control embryo incubated at 24°C/24°C AP/VP (left) and embryo incubated in chamber with temperature gradient 13°C/24°C AP/VP (right). Imaged from 18:04 hpf to 21:54 hpf. View of animal pole surface. Images were collected every 5 minutes; display rate is 6 frames per sec (1 sec of video = 30 min real time). Scale bar 0.2 mm.

Figure S1. **An Embryo Temperature Controller Enables Spatial Control of Cell Division Speed and Blastula Cell Organization**

(**A**) Developmental stages and timing of *Xenopus* embryos growing at different temperatures, converted from data on Xenbase from Khokha et al., Developmental Dynamics, 2002.

**(B)** Schematics of the timing of temperature gradient treatment.

**(C)** Schematics of the chambers of temperature gradient treatment with dimensions.

**(D)** Representative image of a temperature gradient chamber, with a peltier (cooling or heating) on the bottom and a thermistor (temperature measurement) inserted on the side of bottom block. Peltier and thermistor are connected to a temperature controller. Only the bottom peltier and thermistor are shown for clarity.

**(E)** Effective temperature inferred from cell density on each of 12 z sections in embryos. Mean +- SD, N=2 embryos for 22°C, N=6 embryos for 13°C/25°C.

**(F)** Average duration of cell cycles of cells on the surface of animal pole at 23°C. Each dot represents average of durations in 4 to 8 cells in one embryo. Mean +- SD. N=3 embryos. Note that cell cycle elongates at $13^{th}$ and $14^{th}$ cell cycle.

**(G)** Cell cycle number of AP surface cells at 23°C after temperature gradient, inferred from the cell density. Mean +- SD. N=3-12 embryos for 24°C/24°C, N=5 embryos for 13°C/24°C.

**(H)** Schematics of temperature treatment applied to control and two opposite directions of temperature gradient. Bottoms are 9 hpf stereomicroscope images of AP and VP to show cell size comparisons and developmental outcomes respectively. Scale bars, 0.5 mm.

Figure S2 (related to Figures 1-3) – **Side-to-side temperature control or cell division speed creates ectopic blastula cell size and ZGA onset gradients**

**(A)** Top-down pattern of ZGA onset in control embryos. Segmented nuclei shown with heatmap of nascent transcription (EU-RNA), from 7-9 hpf. Nascent transcription in detectable first in AP cells and later in equatorial regions and finally in VP cells.

**(B)** Schematics of embryos undergoing normal temperature or side-to-side temperature gradient. Right: expected spatial patterns of genome activation.

**(C)** Stereomicroscope images of animal pole of 22°C|22°C control and 15°C|22°C embryo at 8.3 hpf. Scale bar, 0.2 mm.

**(D)** Nuclei position plots color-coded with cell size (diameter). Left: a 22°C|22°C control embryo. Right: a 15°C|22°C embryo. Top: top views of top 1/4 sections of embryos. Bottom: side views of whole embryos.

**(E)** Nuclei position plots color-coded with cell size (top) or EU amount (bottom). Center 1/6 sections. Left: a 22°C|22°C control embryo. Right: a 15°C|22°C embryo.

**(F)** Percentages of transcription active cells in each bin of cell sizes (every 2 μm, 15 bins from 30 μm to 60 μm). Mean +- SD, (N=2 embryos for 22°C, N=4 embryos for 15°C|22°C).

Figure S3, related to Figure 3

**Transcription delay in inverted embryos induces apoptosis.**

**(A)** Percentages of embryo deaths in different groups of temperature treatment. N=8 experiments for each different temperature. Mean +- SD. N=12 or 24 embryos per experiment. Multiple paired parametric t-tests, P=0.0056 or 0.0019.

**(B)** Representative 24°C/24°C and 13°C/24°C embryos at 24 hpf. Confocal slices showing DNA staining (magenta) and cleaved caspase 3 staining (green) with 3x zoom-in views and schematics of early neurula (24°C/24°C) and apoptotic embryos (13°C/24°C) in which apoptotic cells mixed with normal cells in equatorial region and released into blastocoel. Scale bars, 80 μm in both views. Drawing of neurula: *Xenopus* illustrations © Natalya Zahn (2022).

**(C)** Representative results of Western blot of β-tubulin or cleaved caspase 3. Embryos were collected at 26.6 hpf after temperature treatment of 24°C/24°C or 13°C/24°C. N=10 embryos pooled in each sample.

Figure S4, related to Figure 4

**Embryo quality control senses ZGA inhibition, and revealing the origin of apoptotic cells**

**(A)** Representative results of Western blot of β-tubulin or cleaved caspase 3. Embryos were collected at 23.2 hpf after TBS buffer or α-amanitin microinjections. N=10 embryos pooled in each sample.

**(B)** Levels of cleaved caspase 3 relative to β-tubulin in Western blot, shown as fold change in α-amanitin-injected embryos relative to TBS-injected embryos. Each dot represents one Western blot. Samples collected from two independent experiments. N=10 embryos pooled in each sample. Mean +- SD. Ratio paired parametric t-test, P<0.0001.

**(C)** Left column: representative stereomicroscope images of embryos at 31 hpf, tailbud stage, with normal ZGA, blocked ZGA by α-amanitin, or blocked ZGA but rescued by caspase inhibitors. Right column: schematics of development outcome of three treatments. Scale bar, 0.5 mm. Drawing of tailbud: *Xenopus* illustrations © Natalya Zahn (2022).

**(D)** Death percentages of each group, only the embryos with blocked transcription showed death percentages of 100%. N=4 to 14 experiments. N=15-30 embryos per experiment. Mean +- SD. Mann-Whitney tests, P<0.0001.

**(E)** Top views of the of embryos undergoing apoptosis after temperature gradient or α-amanitin injections, with apoptotic cells released from blastocoel to extra-embryonic space. Arrows pointed to the direction in which white apoptotic cells are floating. Brightness and saturation were adjusted in images of the top row. Scale bar, 0.5 mm.

**(F)** Schematics showing 1-cell α-amanitin injection followed by dye injections at the 32-cell stage. B1-labeled but not D1-labeled cells are expected to be apoptotic and dissociate. Arrows showing the apoptotic cells breaking into perivitelline space. Lineage tracing maps of embryos at 32-cell stage, stage 8, and stage 10.5 are adopted from Xenbase.

**(G)** Top views of B1-labeled embryos with normal or blocked ZGA. Brightfield, GFP channel, and the merged views are shown. At 5.5 hpf, embryos showed fluorescent cells on the edge of the animal cap. At 12 hpf, B1-labeled α-amanitin-injected embryos showed many fluorescent cells dissociating from embryos (numbers of embryos shown on top right). TBS-injected embryos did not show dissociated cells. Scale bar, 0.5 mm.

**(H)** Top views of D1-labeled embryos with normal or blocked ZGA. At 5.5 hpf, embryos showed fluorescent cells on the edge of the vegetal side. At 12 hpf, D1-labeled α-amanitin-injected embryos did not show fluorescent cells dissociating from embryos. TBS-injected embryos did not show dissociated cells. Scale bar, 0.5 mm.

**(I)** At 12 hpf, B1-labeled α-amanitin-injected embryos showed significantly higher percentages of fluorescent apoptotic cell release compared to D1-labeled α-amanitin-injected embryos. Mean +- SD. N=2 or 4 experiments. N=13 embryos per experiment on average. Unpaired parametric t-test, P=0.0005.

Figure S4 continued

**(J)** At 12.5 hpf, B1-labeled α-amanitin-injected embryos showed many fluorescent cells dissociating from embryos (arrows), when blastocoels were cut open. Scale bar, 0.5 mm.

**(K)** At 12.5 hpf, D1-labeled α-amanitin-injected embryos showed very few fluorescent cells dissociating from embryos, when blastocoels were cut open. Scale bar, 0.5 mm.

**(L)** Schematics showing 1-cell α-amanitin injection followed by dye injections at the 32-cell stage. Some of the C1-labeled and C4-labeled cells are expected to be apoptotic and dissociate. Lineage tracing maps of embryos at 32-cell stage, stage 8, and stage 10.5 are adopted from Xenbase.

**(M)** Top views of C1-labeled embryos with normal or blocked ZGA. At 5.5 hpf, embryos showed fluorescent cells on the dorsal equatorial region. At 12 hpf, C1-labeled α-amanitin-injected embryos showed fluorescent cells dissociating from embryos. TBS-injected embryos did not show dissociated cells. Scale bar, 0.5 mm.

**(N)** Bottom views of C4-labeled embryos with normal or blocked ZGA. At 5.5 hpf, embryos showed fluorescent cells on the ventral equatorial region. At 12 hpf, C4-labeled α-amanitin-injected embryos showed fluorescent cells dissociating from embryos. TBS-injected embryos did not show dissociated cells. Scale bar, 0.5 mm.

**(O)** At 12.5 hpf, C1-labeled α-amanitin-injected embryos showed many fluorescent cells dissociating from embryos (arrows), when blastocoels were cut open. Scale bar, 0.5 mm.

**(P)** At 12.5 hpf, C4-labeled α-amanitin-injected embryos showed many fluorescent cells dissociating from embryos, when blastocoels were cut open. Scale bar, 0.5 mm.

**(Q)** At 12 hpf, B1, C1, or C4-labeled α-amanitin-injected embryos showed higher ratios of fluorescent apoptotic cell release compared to D1-labeled α-amanitin-injected embryos. Mean +- SD. N=1, 2, or 4 experiments. N=12 embryos per experiment on averagee

### Figure S1

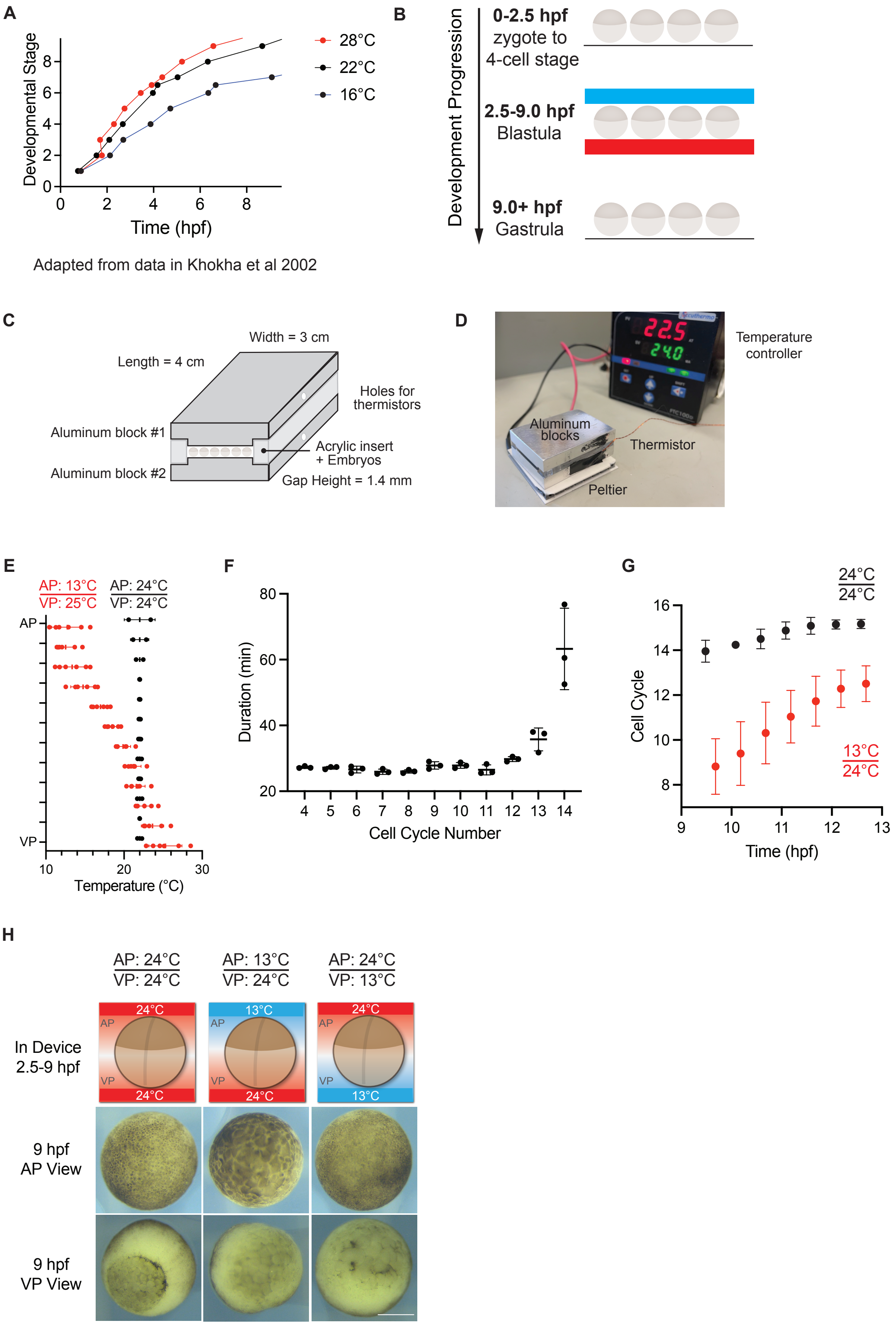

A
Developmental Stage
Time (hpf)
28°C
22°C
16°C
Adapted from data in Khokha et al 2002
B
Development Progression
0-2.5 hpf
zygote to
4-cell stage
2.5-9.0 hpf
Blastula
9.0+ hpf
Gastrula
C
Width = 3 cm
Length = 4 cm
Holes for
thermistors
Aluminum block #1
Aluminum block #2
Acrylic insert
+ Embryos
Gap Height = 1.4 mm
D
Temperature
controller
Aluminum
blocks
Thermistor
Peltier
E
AP: 13°C
VP: 25°C
AP: 24°C
VP: 24°C
AP
VP
Temperature (°C)
F
Duration (min)
Cell Cycle Number
G
24°C
24°C
Cell Cycle
13°C
24°C
Time (hpf)
H
AP: 24°C
VP: 24°C
AP: 13°C
VP: 24°C
AP: 24°C
VP: 13°C
In Device
2.5-9 hpf
9 hpf
AP View
9 hpf
VP View

### Figure S2

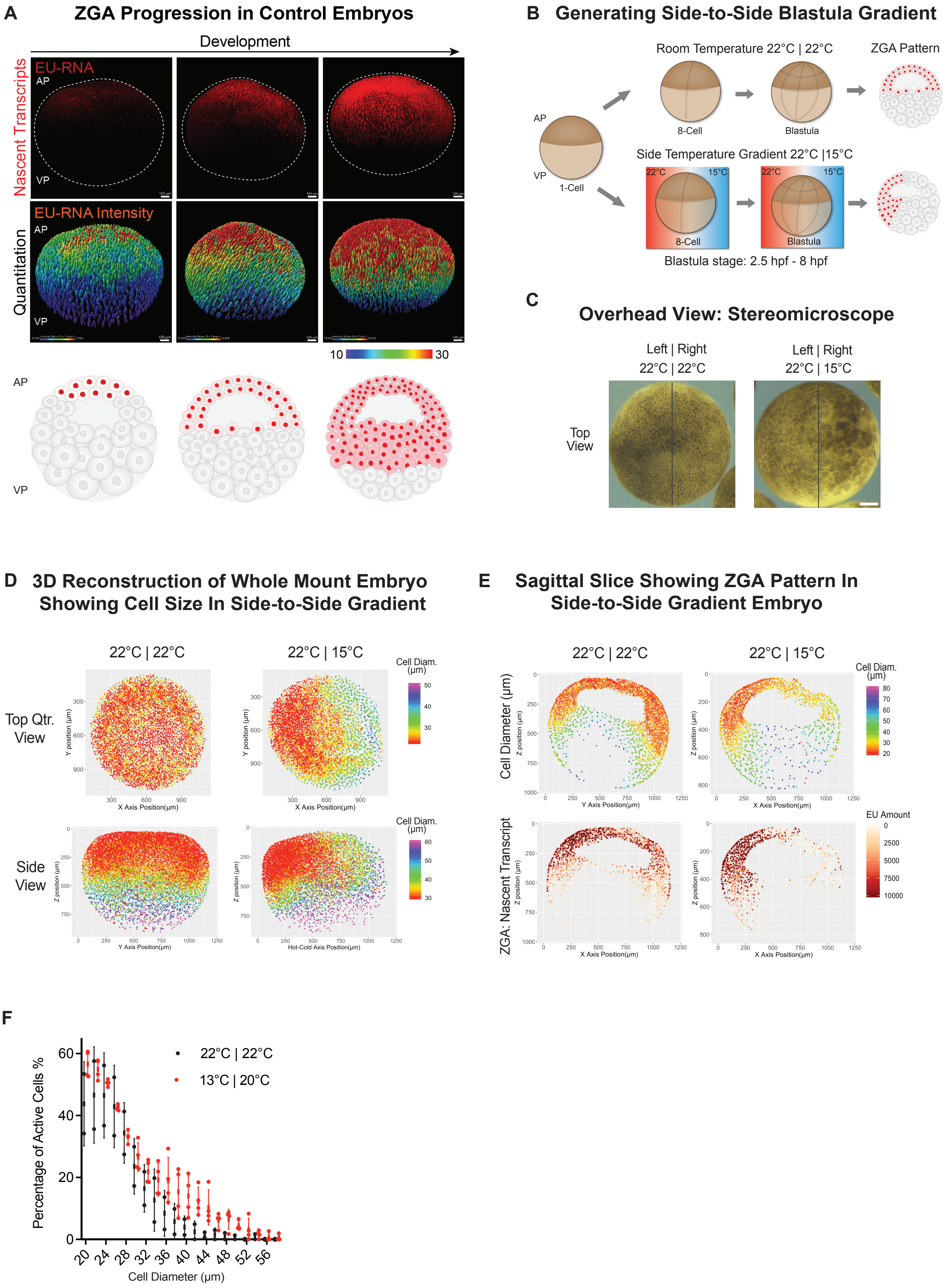

A
ZGA Progression in Control Embryos
Development
Nascent Transcripts
EU-RNA
AP
VP
Quantitation
EU-RNA Intensity
10
30
B
Generating Side-to-Side Blastula Gradient
Room Temperature 22°C | 22°C
ZGA Pattern
8-Cell
Blastula
1-Cell
Side Temperature Gradient 22°C |15°C
22°C
15°C
Blastula stage: 2.5 hpf - 8 hpf
C
Overhead View: Stereomicroscope
Left | Right
22°C | 22°C
22°C | 15°C
Top View
D
3D Reconstruction of Whole Mount Embryo Showing Cell Size In Side-to-Side Gradient
22°C | 22°C
22°C | 15°C
Cell Diam. (µm)
Top Qtr. View
Side View
Y position (µm)
Z position (µm)
X Axis Position(µm)
Y Axis Position(µm)
Hot-Cold Axis Position(µm)
E
Sagittal Slice Showing ZGA Pattern In Side-to-Side Gradient Embryo
Cell Diameter (µm)
ZGA: Nascent Transcript
EU Amount
F
Percentage of Active Cells %
Cell Diameter (µm)
22°C | 22°C
13°C | 20°C

Figure S3

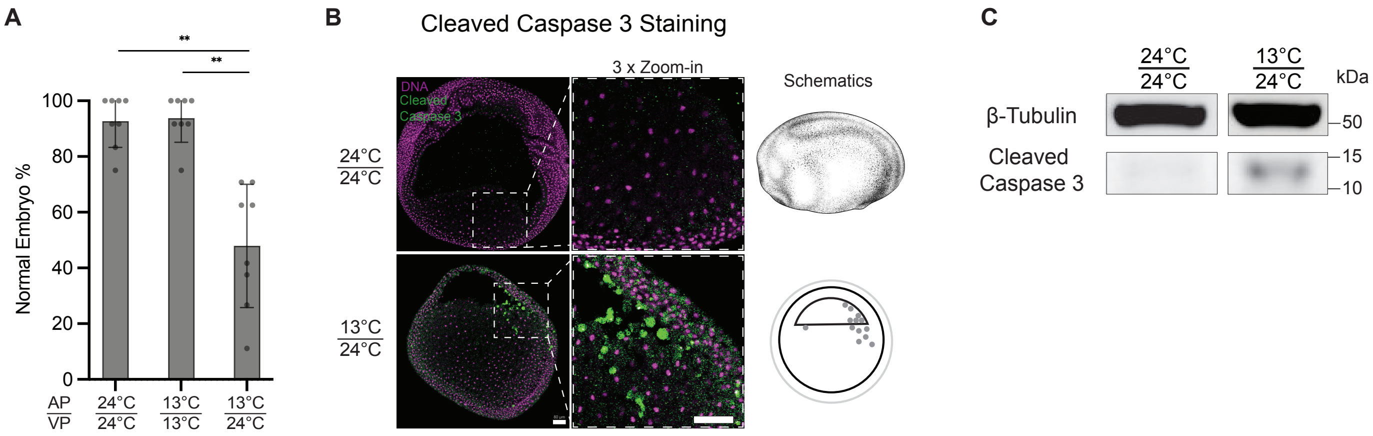
A
**
**
Normal Embryo %
100
80
60
40
20
0
AP/VP
24°C/24°C
13°C/13°C
13°C/24°C
B
Cleaved Caspase 3 Staining
3 x Zoom-in
Schematics
DNA
Cleaved
Caspase 3
24°C/24°C
13°C/24°C
C
24°C/24°C
13°C/24°C
kDa
β-Tubulin
50
Cleaved
Caspase 3
15
10

### Figure S4

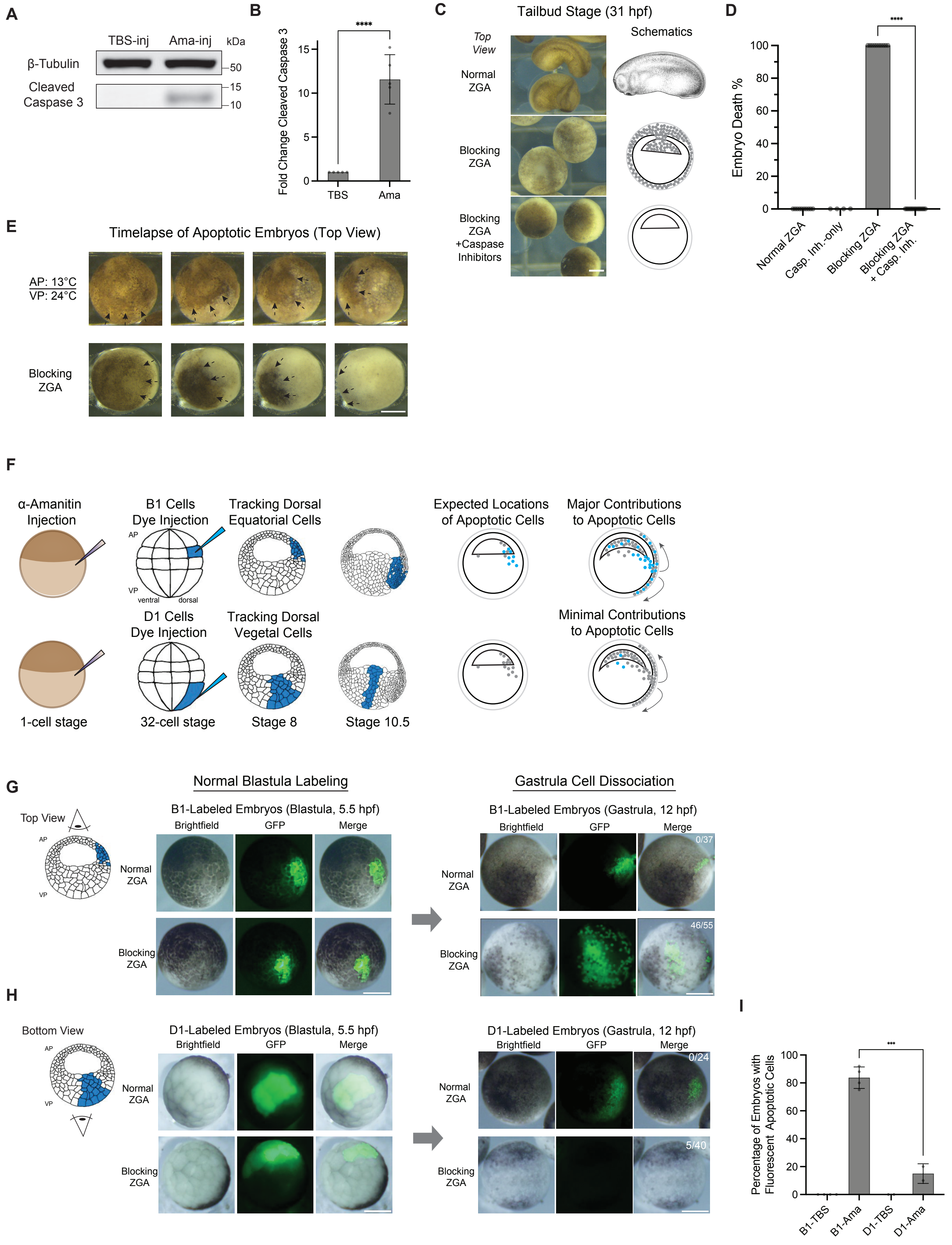

A
TBS-inj
Ama-inj
kDa
β-Tubulin
Cleaved Caspase 3
50
15
10
B
Fold Change Cleaved Caspase 3
****
TBS
Ama
C
Tailbud Stage (31 hpf)
Top View
Schematics
Normal ZGA
Blocking ZGA
Blocking ZGA +Caspase Inhibitors
D
Embryo Death %
****
Normal ZGA
Casp. Inh.-only
Blocking ZGA
Blocking ZGA + Casp. Inh.
E
Timelapse of Apoptotic Embryos (Top View)
AP: 13°C
VP: 24°C
Blocking ZGA
F
α-Amanitin Injection
B1 Cells Dye Injection
Tracking Dorsal Equatorial Cells
Expected Locations of Apoptotic Cells
Major Contributions to Apoptotic Cells
AP
VP
ventral
dorsal
D1 Cells Dye Injection
Tracking Dorsal Vegetal Cells
Minimal Contributions to Apoptotic Cells
1-cell stage
32-cell stage
Stage 8
Stage 10.5
G
Normal Blastula Labeling
Gastrula Cell Dissociation
Top View
B1-Labeled Embryos (Blastula, 5.5 hpf)
B1-Labeled Embryos (Gastrula, 12 hpf)
Brightfield
GFP
Merge
Normal ZGA
Blocking ZGA
0/37
46/55
H
Bottom View
D1-Labeled Embryos (Blastula, 5.5 hpf)
D1-Labeled Embryos (Gastrula, 12 hpf)
0/24
5/40
I
Percentage of Embryos with Fluorescent Apoptotic Cells
***
B1-TBS
B1-Ama
D1-TBS
D1-Ama

### Figure S4 continued

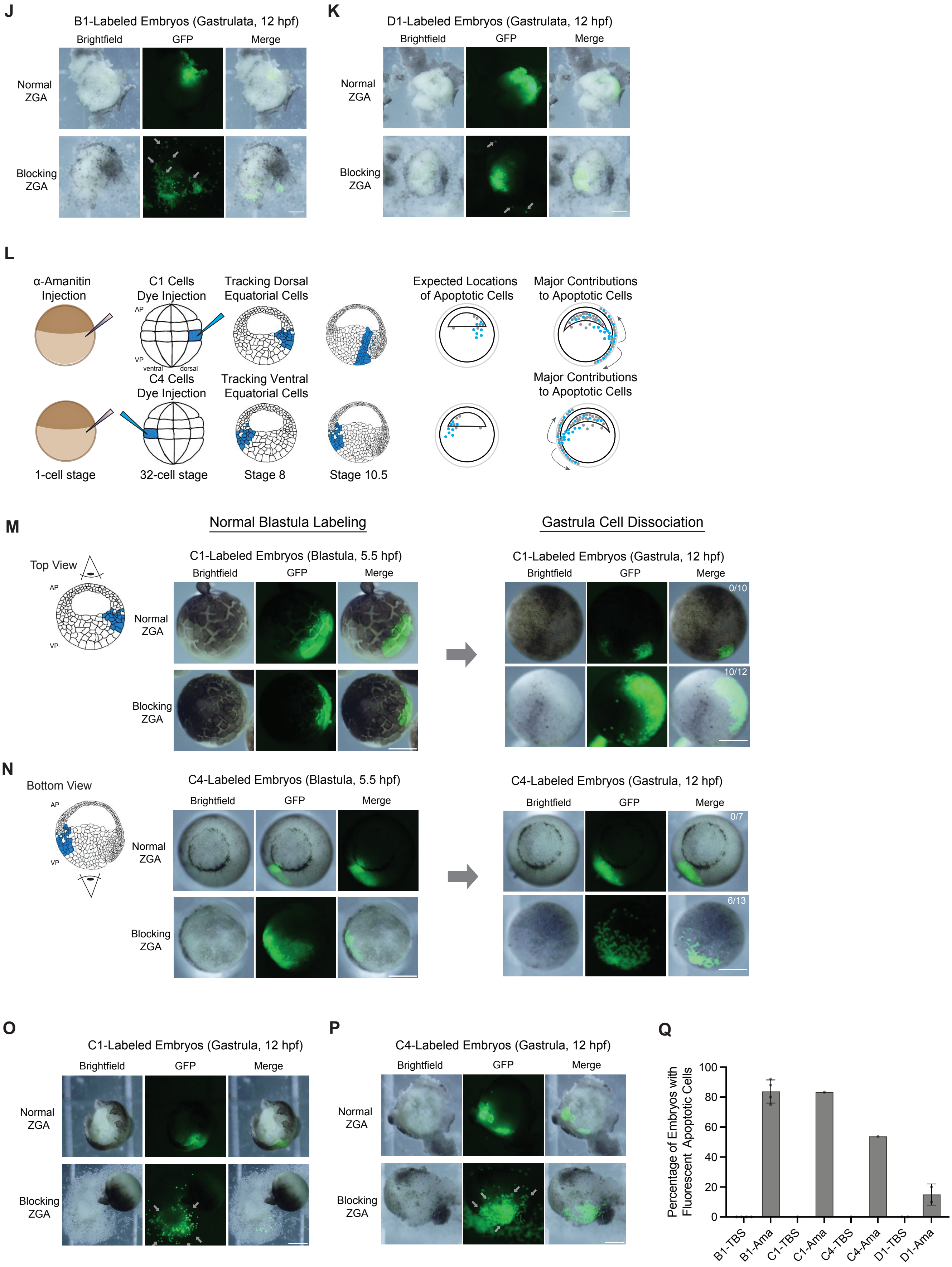
